# Microscopic interactions control a structural transition in active mixtures of microtubules and molecular motors

**DOI:** 10.1101/2023.01.04.522797

**Authors:** Bibi Najma, Aparna Baskaran, Peter J. Foster, Guillaume Duclos

## Abstract

Microtubules and molecular motors are essential components of the cellular cytoskeleton, driving fundamental processes *in vivo,* including chromosome segregation and cargo transport. When reconstituted *in vitro*, these cytoskeletal proteins serve as energy-consuming building blocks to study the self-organization of active matter. Cytoskeletal active gels display rich emergent dynamics, including extensile flows, locally contractile asters, and bulk contraction. However, how the protein-protein interaction kinetics set their contractile or extensile nature is unclear. Here, we explore the origin of the transition from extensile bundles to contractile asters in a minimal reconstituted system composed of stabilized microtubules, depletant, ATP, and clusters of kinesin-1 motors. We show that the microtubule binding and unbinding kinetics of highly processive motor clusters set their ability to end-accumulate, which can drive polarity sorting of the microtubules and aster formation. We further demonstrate that the microscopic time scale of end-accumulation sets the emergent time scale of aster formation. Finally, we show that biochemical regulation is insufficient to explain fully the transition as generic aligning interactions through depletion, crosslinking, or excluded volume interactions can drive bundle formation, despite the presence of end-accumulating motors. The extensile-to-contractile transition is well captured by a simple self-assembly model where nematic and polar aligning interactions compete to form either bundles or asters. Starting from a five-dimensional organization phase space, we identify a single control parameter given by the ratio of the different component concentrations that dictates the material-scale organization. Overall, this work shows that the interplay of biochemical and mechanical tuning at the microscopic level controls the robust self-organization of active cytoskeletal materials.

**Significance statement:** Self-organization in living cells is often driven by energy-consuming motor proteins that push and pull on a network of cytoskeletal filaments. However, it is unclear how to connect the emergent structure and dynamics of reconstituted cytoskeletal materials to the kinetics and mechanics of their microscopic building blocks. Here, we systematically correlate bulk structure with asymmetry of the motor distribution along single filaments to explain the transition from extensile bundles to contractile asters in active networks of stabilized microtubules crosslinked by motor proteins. We combine experiments and scaling arguments to identify a single number that predicts how the system will self-organize. This work shows that biochemical and mechanical interactions compete to set the emergent structure of active biomimetic gels.

## Introduction

Active materials are far-from-equilibrium materials composed of energy-consuming building blocks [1]. They self-organize into spontaneously moving structures larger than their microscopic components, and their material-scale mechanics emerge from non-equilibrium interactions between active units. In thermodynamically driven self-assembly, the emergent structures are set by the interaction strengths of the various molecules [2]. In active materials, pattern formation often results from dynamical processes whose characteristic time scale sets the emergent structure and dynamics [3].

Many examples of *in vitro* cytoskeletal materials self-organize into distinct states, which reflect the richness of the microscopic interactions between various molecular motors and cytoskeletal filaments. In particular, these materials often display bulk extensile [4-6] or contractile and aster-forming behaviors [7-18]. Extensile dynamics have been associated with active systems organized into mesoscopic polymer bundles, frequently through the addition of a depletion agent. Such extensile flows have been reported for both microtubule/kinesin [4,6,18] and actomyosin [19,20] based materials. Theoretical work has suggested that extensile stresses could arise from the polarity sorting of filaments [21] or the mechanical properties of the motors themselves [22,23]. In the absence of depletant, both microtubule and actin-based active gels have been found to undergo bulk contraction. However, the mechanisms that lead to extensile or contractile dynamics might differ as actin and microtubules have very different mechanical properties. Actomyosin contraction is also fundamentally rooted in symmetry-breaking at the filament level, either because of the non-linear mechanical properties of actin filaments [11,24-26], end-accumulation of myosin motors [16,27], or through the asymmetric distribution of crosslinkers or motor proteins [28]. Microtubule bulk contraction has been argued to stem from contractile active stresses generated by the clustering of microtubule ends by end-accumulating motors [7,12,29,30].

While dyneins and many kinesins are known to end-accumulate [31-34], many motors - including dimeric kinesin-1 – have been argued not to under physiological conditions, even on stabilized microtubules [32,35,36]. Indeed, theory and experiments demonstrated that the processivity and binding and unbinding kinetics play a crucial role in determining whether molecular motors will end-accumulate on isolated stabilized microtubules [32,37]. Furthermore, the kinetics of the motor binding and unbinding depends on several factors, including salt concentration and the concentrations of motors and ATP [38,39]. Finally, as processivity increases when motors can bind to multiple sites along a single filament [40], multivalency of the motors is important to consider as many motors form clusters, either naturally or by design [4,7,19,41]. Thus, whether or not a motor will end-accumulate is determined in part by the motor’s environment.

On the theoretical front, simple theories and simulations have addressed the origin of contractility in crosslinked networks composed of rigid or semi-flexible polymers and molecular motors [25,30,42], highlighting the importance of having passive crosslinkers that create an elastically percolated network. For end-accumulating motors, experiments and computer simulations showed that the competition between motor crosslinking either the ends or the sides of microtubules set the extensile or contractile dynamics of the active network [43]. However, despite these advances, the rational design of a simple and programmable biomimetic material that could exert both extensile and contractile stresses is still challenging. Recent *in vitro* experiments and simulations show that microtubule networks composed of a mixture of motors with opposite directionality can display either extensile or contractile dynamics depending on the relative concentration of each motor protein [43,44]. In the presence of a single type of end-accumulating motor, competition between microtubule growth rate and motor speed dictates if bulk dynamics is extensile or contractile [43].

Most reconstituted active systems are composed of stabilized microtubules and a single type of motor, and were thought to display a single bulk organization - either contractile or extensile [4,7,12]. However, two recent examples, with clusters of kinesin-1 motors [18,45] and end-accumulating kinesin-4 motors [34], show that networks of stabilized microtubules crosslinked by a single type of motor proteins can display both extensile and contractile dynamics. The underlying physical or biochemical mechanisms that lead to this phenomenon are unclear. In particular, the role of motor-microtubule interaction kinetics in determining the emergent structure of the active microtubule-kinesin gel is unknown. There is, therefore, a need for a careful study of the simplest system possible, with stabilized microtubules and a single motor type, to uncover the microscopic origin of the extensile-to-contractile transition.

Here, we assembled an active gel composed of stabilized microtubules, a bundler, and kinesin-1 motor clusters. We systematically combined bulk experiments with direct observation at the single filament level to study how emergent bulk dynamics is related to the distribution of processive motor clusters along microtubules. Extensile flows emerged when the motors were uniformly distributed and the active gel contracted when motors end-accumulated. We show that end-accumulation and contraction can be induced by either increasing motor cluster concentration or decreasing ATP concentration, which is consistent with increasing the lattice-binding rate and decreasing the lattice-dissociation rate of the motor-microtubule binding kinetics. Our study goes beyond steady-state dynamics, showing that the microscopic timescale of end-accumulation and the macroscopic timescale for aster formation are correlated. Finally, increasing nematic alignment by adding either a non-specific depletant, a microtubule-specific crosslinker, or colloidal rods, triggers the reversed transition from asters to bundles without impacting motor cluster end-accumulation at the single filament level. A simple self-assembly model demonstrates that the bundle-aster phase boundary results from the competition between polar sorting and nematic alignment. This model allows us to predict the emergent structure based on a single control parameter given by the ratio of the various component concentrations.

## Results

Our minimal *in vitro* system is composed of microtubules and multivalent clusters of molecular motors (**Fig. 1A**) [4,46]. GMPCPP-stabilized microtubules are bundled by a non-specific depletant (20kDa PEG, **Fig. 1B**). Motor clusters are composed of tetravalent streptavidin and biotinylated truncated kinesin-1 motors (consisting of the first 401 amino acids of kinesin-1) that spontaneously dimerize to form a processive double-headed motor (**Fig. 1C**). The exact valency and size of these clusters are unknown. These motor clusters can simultaneously bind to multiple microtubules, taking steps upon hydrolysis of adenosine 5ʹ-triphosphate (ATP). This crosslinking and walking motion results in the sliding of adjacent antiparallel microtubules. In this configuration, a microtubule bundle continuously elongates over time and exerts dipolar extensile stresses [4,6,47] (**Fig. 1B**). Microtubule bundles form a continuously reconfiguring three-dimensional active gel that powers autonomous extensile flows [4,48] (**Fig. 1D, Video S1**). In this extensile phase, dual-color imaging of microtubules and motor clusters revealed uniform distribution of the motors along the microtubule bundles (**Fig. 1G, H, Fig. S1A-C**).

**Figure 1:**
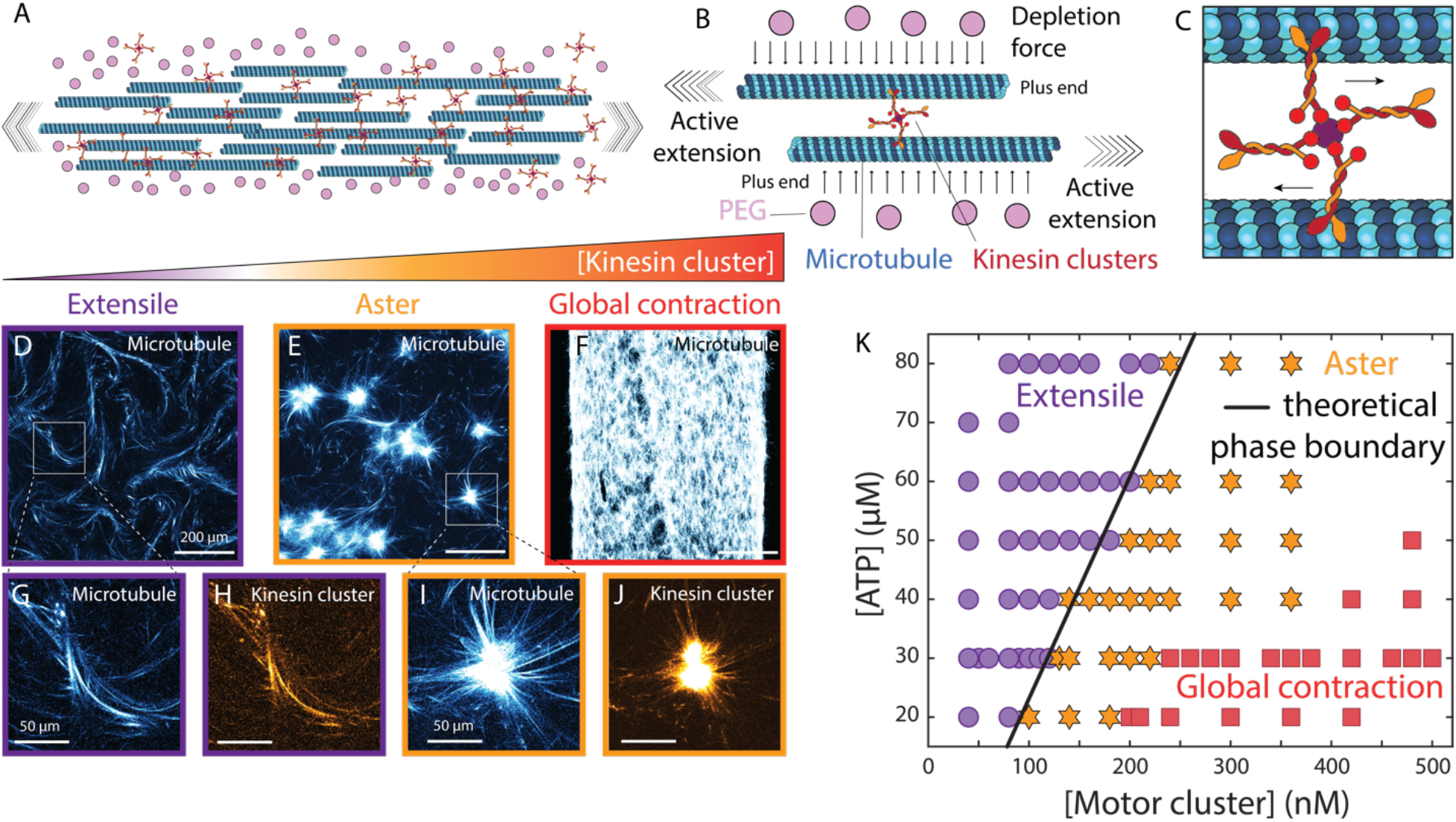
Multivalent clusters of Kinesin-1 molecular motor drive microtubule gels into extensile, aster, or globally contractile phases. A) Schematic of an extensile bundle of stabilized microtubules. B) Close up on anti-parallel microtubules bundled by a non-specific depletant (PEG). C) Close up on the multivalent clusters of kinesin-1 motors. Motors are biotinylated and dimerize spontaneously. Clusters are formed by tetravalent streptavidin. Kinesin-1 motors walk toward the plus end of the microtubules. As a result, anti-parallel microtubules slide apart when both are bound to a motor cluster. D) Fluorescent picture of the microtubule gel in the extensile phase ([motor clusters]= 60 nM). Increasing the cluster concentration first leads to the formation of E) an aster phase that locally contracts ([motor clusters]= 120 nM) and then F) a globally contractile phase ([motor clusters]= 480 nM). G and H show close-ups of fluorescently labeled tubulin and motor clusters for an extensile bundle. I and J show close-ups of fluorescently labeled tubulin and motor clusters for an aster. K) ATP-motor cluster phase diagram: the critical concentration of motors needed to form asters and contract depends on the ATP concentration. Purple disks represent extensile bundles, orange stars are asters, and red squares are globally contractile gels. Each data point is an independent experiment. D-J were taken with a confocal microscope. Scale bars in D-F are 200 µm. Scale bars in G-J are 50 µm. In D-K, [ATP]= 30 µM, [PEG]=0.6% vol/vol, [tubulin]=1.33mg/mL.

Assembling active gels with increasing motor cluster concentrations led to a radical change in the emergent structure and dynamics of the active gels: above a critical motor cluster concentration, the active network transitioned from forming an extensile phase into forming a contractile phase, where large microtubule asters coexist with an extensile background (**Fig. 1E, Video S2**). Dual-color imaging of microtubules and kinesin-1 clusters revealed that the aster cores were enriched with motor clusters (**Fig. 1I, J, Fig. S1D-F**), a generic feature of both actin-based [16,27] and microtubule-based asters, both *in vitro* [18,34,43,49] and in cellular extracts [12,50-52]. Further increasing the motor cluster concentration led to a second transition from a locally contractile aster phase to a globally contractile phase (**Fig. 1F, Video S3**).

We further investigated how the binding and unbinding rates of kinesin onto microtubules control the macroscopic organization of the network by changing the concentrations of ATP and motor clusters while keeping the microtubules’ length and the concentrations of microtubules and PEG constant. The resulting phase diagram reveals that the critical motor cluster concentration that triggers aster formation depends on the ATP concentration (**Fig. 1K**). As the asters displayed enrichment of motors in their core (**Fig. 1J, Fig. S1D-F)**, a hallmark of asymmetric motor distribution, we hypothesized that modulating the ATP and kinesin concentrations changes the degree of motor end-accumulation, and hence the system’s propensity towards contractility.

To test this hypothesis, we looked at the distribution of fluorescent motor clusters along isolated microtubules (**Fig. 2A, Supplementary Materials, section 2a**). We observed that motor clusters were uniformly distributed along microtubules in conditions where the gel was extensile (**Fig. 2C, F**). However, when the concentration of ATP was decreased (**Fig. 2B, D**) or the concentration of motor clusters was increased (**Fig. 2G,H**), motor clusters formed dense aggregates on one end of the microtubules and the corresponding bulk dynamics were contractile. We confirmed these trends by quantifying the fraction of microtubules with motor cluster caps for a wide range of ATP and motor cluster concentrations (**Supplementary Materials, section 4a**). Decreasing ATP concentration or increasing motor cluster concentration led to an increase in the fraction of microtubules with motor caps (**Fig. 2E, I**). Interestingly, we report that the probability of having a cap increased with microtubule length (**Fig. S2E**). Hence, an antenna model may underlie the cap formation process: more motors can be captured by longer microtubules, resulting in a higher probability to form a motor cap [32,53]. We tested the impact of microtubule length on the macroscale organization by annealing microtubules while fixing the tubulin concentration: microtubules got longer while their number density decreased. This led to a transition from extensile bundles to contractile asters (**Fig. S2A-D**). Decreasing microtubule density while keeping their average length constant also led to a transition from extensile to contractile dynamics (**Fig. S3A**).

**Figure 2:**
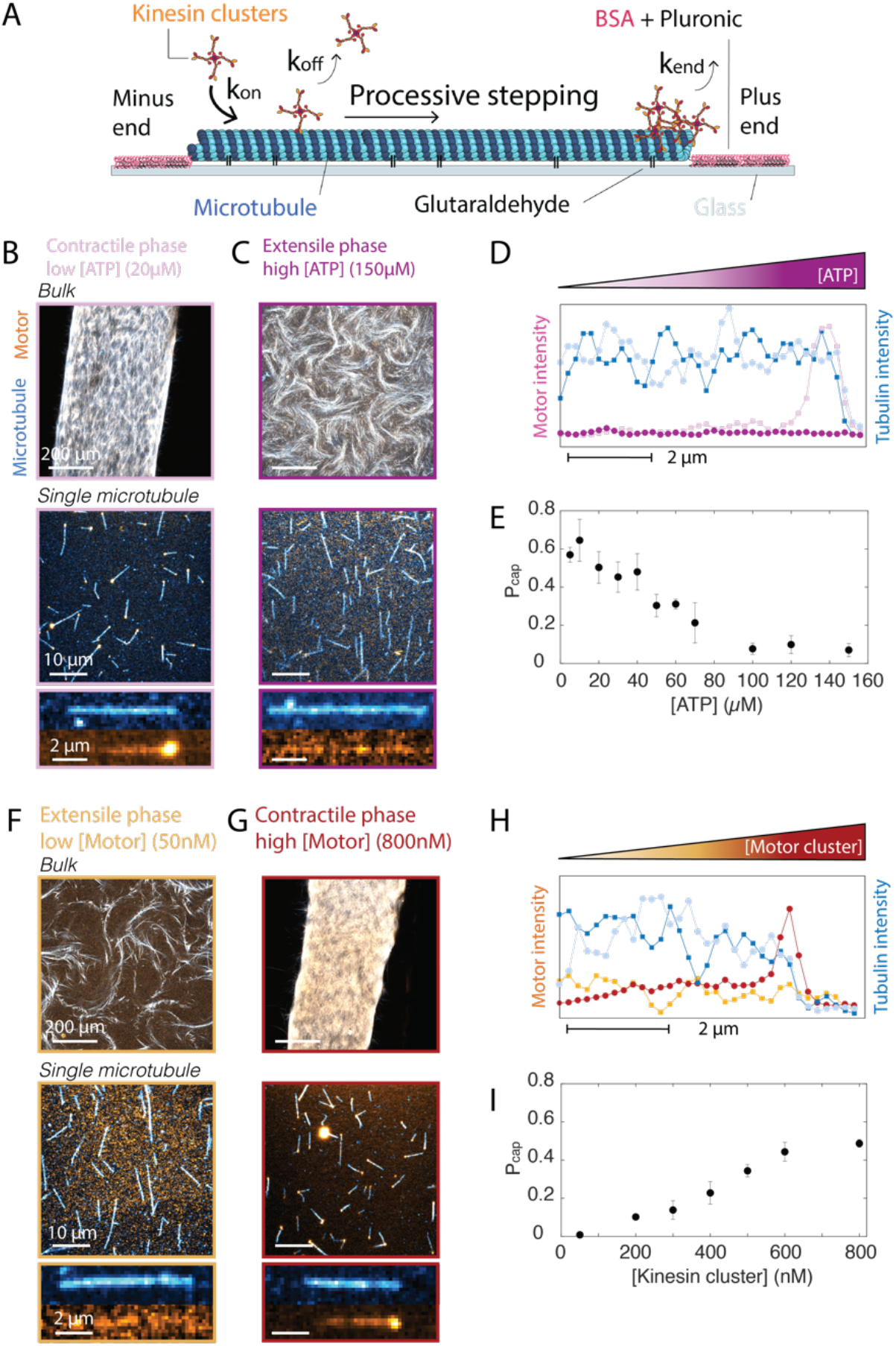
Kinetics of kinesin cluster-microtubule binding and unbinding lead to motor cluster accumulation at the end of a microtubule. A) Schematic representation of the microscopic interaction between a microtubule and multivalent motor clusters. B-C) Fluorescent pictures of bulk active gels and corresponding fluorescent pictures of isolated microtubules bound to a glass surface at low and high ATP concentrations. D) Line scans of the fluorescent intensity along a single microtubule for the pictures shown in B and C. Microtubule profiles are in cyan, and motor cluster profiles are in magenta. E) Fraction of microtubules with a motor cluster cap for increasing ATP concentrations. F-G) Fluorescent pictures of bulk active gels and corresponding fluorescent pictures of isolated microtubules bound to a glass surface at low and high motor cluster concentrations. H) Line scans of the fluorescent intensity along a single microtubule for the pictures shown in F and G. Microtubule profiles are in cyan, and motor cluster profiles are in orange. I) Fraction of microtubules with a motor cluster cap for increasing motor cluster concentrations. In B-C and F-G, Tubulin is labeled in cyan, motor clusters are in orange. For B-E, [motor cluster]=480 nM; for F-I, [ATP]=30 µM.

Inspired by experiments and theory describing the mechanisms that underlie end-accumulation of highly processive Kip3 motors on microtubules [32], we hypothesized that the processivity of the motor clusters is a key feature that enables the formation of caps and the assembly of contractile asters. We measured the run length of K401 motor clusters under conditions where they end-accumulate and found an average run length of 5.7 ± 2.6 µ*m* (N=216 runs, **Fig. S4A-B**), while the mode of the microtubule length distribution was around 1.6 µm (N>6000 microtubules, **Fig. S4C**). To further confirm the importance of high processivity, we increased the dissociation rate of the motor clusters by increasing the salt concentration. We observed a transition from contractile to extensile bulk dynamics (**Fig. S5A, B, E, F**). Additionally, motor clusters were found to not end-accumulate when salt concentration was increased, consistently with a higher dissociation rate (**Fig. S5C, D, G, H**).

To test non-processive clusters, we leveraged a recently developed light-dimerizable K365 kinesin-1 motor protein (first 365 amino acids of Kinesin-1) [54]. Contrary to the processive dimeric K401 protein, this monomeric derivative of kinesin-1 is non-processive: motors detach from microtubules after each step [55]. Each motor presents either an iLid or a micro tag which dimerize when exposed to blue light [15,56] (**Fig. S6A**). Native gel experiments demonstrated that equimolar mixtures of these two motors do not spontaneously dimerize when not exposed to blue light (**Fig. S6B-C**). Further, as these motor clusters do not contain streptavidin linkers, the valency of the clusters is limited by design to two single-headed non-processive motors. Here, we performed bulk experiments in the absence of depletant to test the ability of these motor dimers to form asters. We combined stabilized microtubules with an equimolar mixture of iLid and micro K365 motors and continuously exposed the sample to blue light while imaging the bulk dynamics. ATP concentrations were varied from 5 µM to 100 µM, and motor concentrations were varied from 100 nM to 500 nM. While all the other kinesin-1 clusters tested assembled microtubules into asters (including light-dimerizable iLid/micro clusters of processive K401 [15]), we never observed the formation of asters with iLid/micro-K365 motor dimers (**Fig. S6D**). Assembling active microtubule gels with iLid/micro dimers of non-processive K365 and a depletant (20 kDa PEG) recovered the extensile dynamics previously reported for this light-activable active gel (**Fig. S6D**) [54].

To further examine the causal link between motor cluster end-accumulation and the formation of contractile asters, we compared the dynamics of both processes (**Fig. 3**). Close to the contractile-to-extensile phase boundary, asters did not appear instantaneously (**Fig. 3A**). Rather, active gels first displayed extensile dynamics before one or more asters nucleated (**Fig. 3A**). We systematically measured the time evolution of aster density and found that increasing ATP concentrations delayed asters nucleation (**Fig. 3B**). Similarly, at the microscopic scale, we found that motor caps formed at slower rates when ATP concentration was increased (**Fig. 3C-D**). We independently measured the time at which the first aster nucleates and the time at which 10% of the microtubules have motor caps under identical buffer conditions. Comparing the two characteristic timescales showed that the dynamics of motor cap and aster formations were correlated (**Fig. 3E**). This analysis provides direct evidence that the microscopic timescale set the emergent timescale.

**Figure 3:**
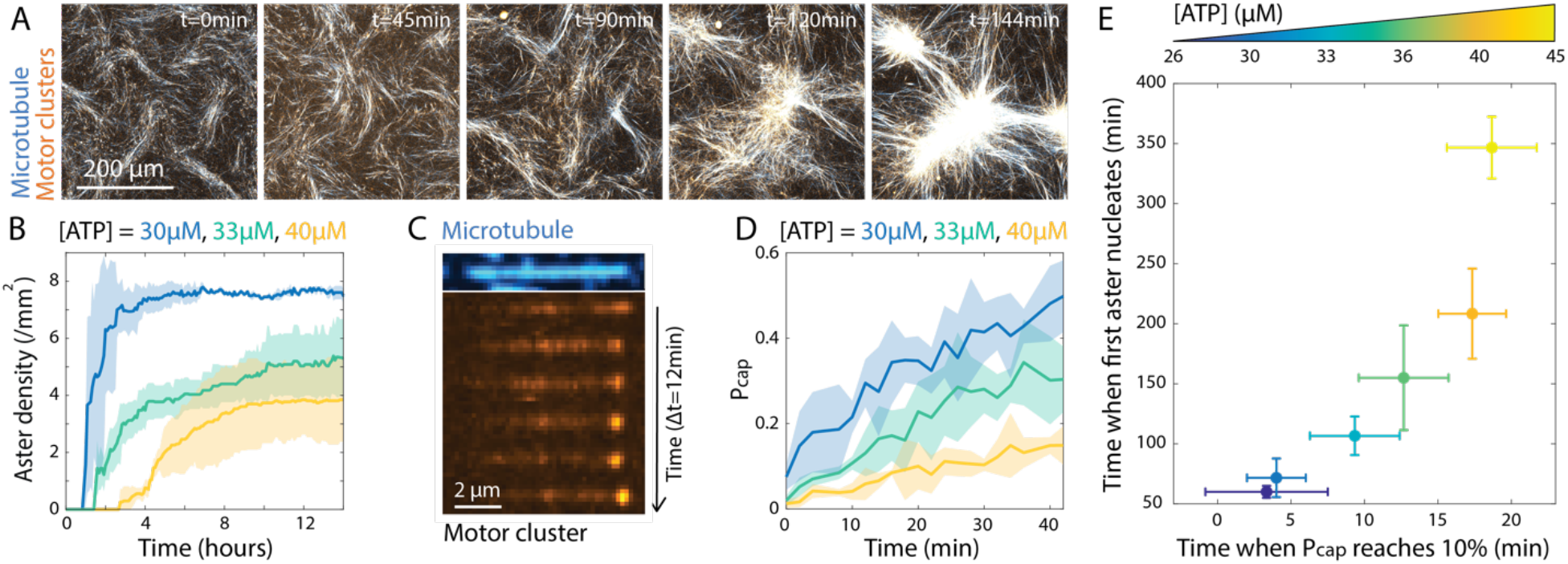
Dynamics of aster formation correlates with dynamics of motor cap formation. A) Time-series of the aster nucleation. Asters nucleate in an extensile background around two hours after the start of the experiment. Tubulin is labeled in cyan, motor clusters are in orange. [ATP]= 33 µM, [motor cluster]=200 nM, [PEG]=0.6% (vol/vol). B) Temporal evolution of aster density for samples with increasing ATP concentrations. Asters nucleate more slowly with more ATP. C) Kymograph (space-time plot) of the formation of a motor cap at the end of a single microtubule. Microtubule is shown in cyan and motor clusters in orange. D) Temporal evolution of P_cap_, the fraction of microtubules with a motor cap, for samples with increasing ATP concentrations. Motor caps form more slowly with more ATP. E) Plot of the time when the first aster nucleates vs time when 10% of the microtubules have a motor cap. Color code shows experiments at various ATP concentrations. A and C were taken with a confocal microscope. Scale bar in A is 200 µm. Scale bar in C is 2 µm. Each experiment in B, D, and E were repeated 3 times independently. Thick lines in B and D and the disks in E show mean values, while shaded areas and error bars show the standard deviation over the 3 replicates.

Taken together, these experiments demonstrate that both the steady-state organization and the dynamics of the self-organization at the macroscopic scale are controlled by the kinetics at the microscopic scale. The microtubule binding and unbinding kinetics of processive motor clusters dictate if and when motors will start to end-accumulate, which is necessary to trigger the transition from extensile bundles to contractile asters. The binding rate of kinesin clusters, k_on_, has been shown to increase when the concentration of motor clusters increases, and the dissociation rate, k_off_, decreases when ATP concentration decreases [39]. Both mechanisms favor the end-accumulation of highly processive motors [32], which, in our case, also triggers end-clustering of the microtubules and aster formation at the macroscopic scale.

However, considering the binding kinetics between motors and microtubules is insufficient to explain fully how kinesin-microtubule networks self-organize in the presence of crowding agents or crosslinkers. Indeed, adding a depletant such as PEG transformed a contractile phase into an extensile phase without impacting the ability of the motor clusters to end-accumulate at the single filament level (**Fig. 4A, B, E, F**). An extensive PEG-motor cluster phase diagram suggests that the contractile-extensile phase boundary results from a competition between the ability to form polar motor caps and the depletion force (**Fig. 4J**). We hypothesized that depletion promotes nematic alignment of microtubules into bundles, while motor cluster caps promote polar sorting and aster formation. We explored how other mechanisms that induce nematic alignment impact the emergent bulk dynamics. We first showed that adding PRC1, a crosslinking protein that binds diffusively to microtubules [57-59], also converted a contractile phase into an extensile phase (**Fig. 4C**). Starting from a globally contractile phase, adding crosslinkers first led to the formation of the aster phase and then an extensile phase (**Fig S7**). Second, we enhanced the nematic alignment of the microtubules by adding passive semiflexible polymers - either filamentous *fd* viruses [60], or DNA-origami rods [61] – to the active sample [62]. These polymers interact sterically with microtubules through excluded volume interaction, enhancing the microtubule alignment and triggering a transition from contractile to extensile dynamics (**Fig. 4D, Fig S8**). Progressively increasing the nematic aligning interaction by adding more colloidal rods converted a globally contractile phase first into an aster phase, then into a fully extensile phase (**Fig. S9A-D**). The critical concentration of colloidal rods needed to trigger the transition increased with motor cluster concentration (**Fig. S9E**). Dual imaging of motor clusters and tubulin at the single microtubule scale revealed that adding PEG, PRC1, or *fd* viruses did not prevent cap formation on isolated microtubules (**Fig. 4E-H**). The fact that increasing depletion, crosslinking, and excluded volume interactions have the same effect on the transition argues that the impact of nematic alignment is generic. To summarize, our results demonstrate that the extensile-contractile transition arises from the competing effects of microtubule end-clustering, which favors the formation of contractile asters, and factors promoting microtubule alignment, which favors the formation of extensile bundles (**Fig. 5A**).

**Figure 4:**
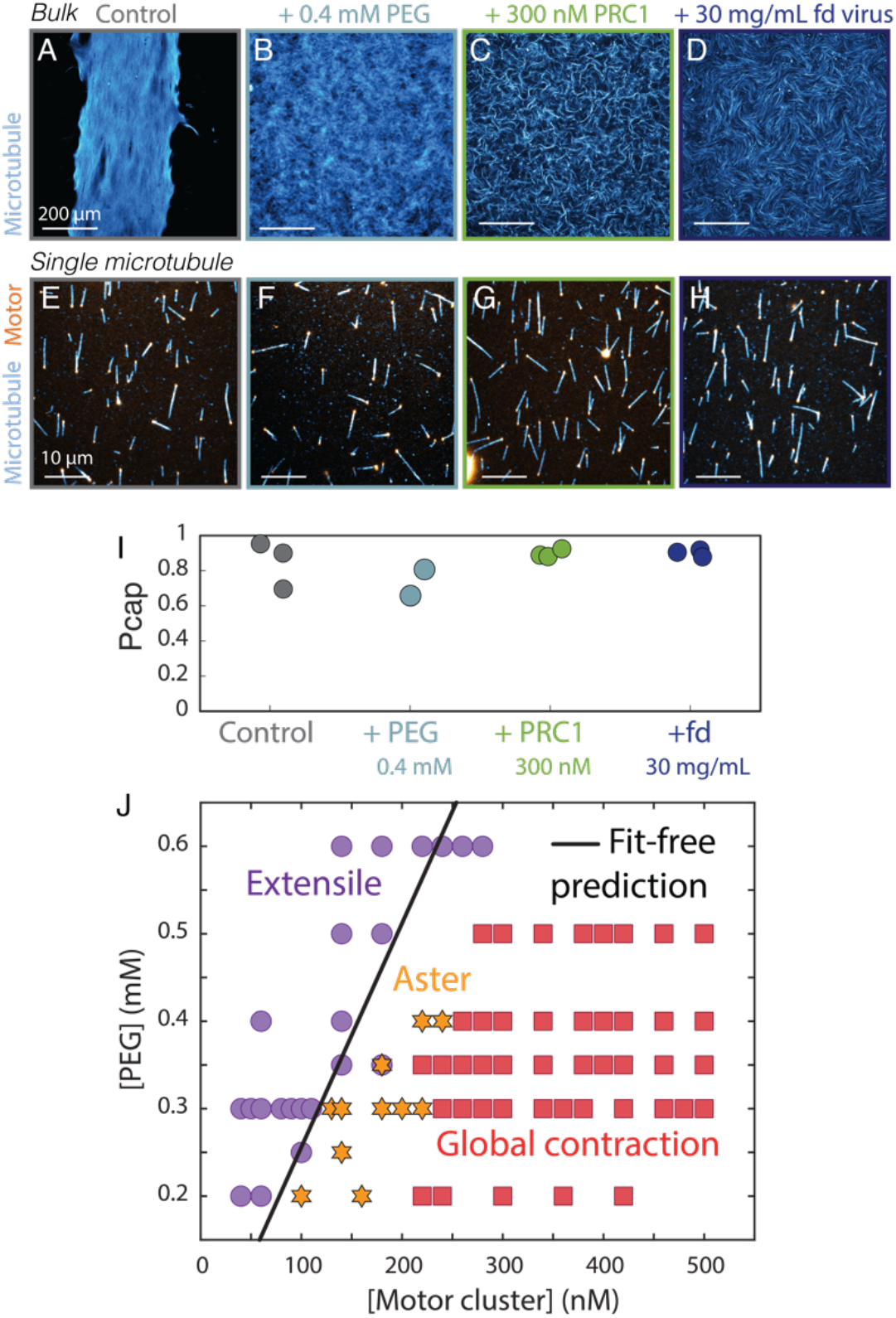
Nematic alignment outcompetes polar sorting to drive a contractile-to-extensile transition, despite the formation of motor caps at the single microtubule level. A) Fluorescent picture of an active crosslinked gel with PRC1 instead of PEG in the globally contractile phase ([ATP]=30 µM, [Motor cluster]=480 nM, [PRC1]=100 nM, [MT]=1.33 mg/mL). Tubulin is labeled in cyan. B) Adding 0.4 mM (0.8 %/% vol) of 20kDa PEG suppressed the global contraction and delayed the appearance of asters from an initially extensile phase by 1 hour. Adding C) 300 nM of PRC1, or D) 30 mg/mL of *fd* virus to increase nematic alignment fully suppresses any contraction over the lifetime of the sample (4 hours). Pictures in A-D were taken 1h after dynamics started. E-H) High-magnification microscopy of isolated microtubules for the same experimental conditions as in A-D. Tubulin is labeled in cyan, motor clusters are in orange. I) Fractions of the microtubules with a motor cap for the same experimental conditions as A-D. J) A PEG-motor cluster phase diagram shows how the ratio of both components sets the extensile or contractile dynamics in active gels. Disks, stars, and squares represent experimental data, while the black line represents the no-fit prediction derived from the self-assembly model and the fitted parameters from the phase diagram in Fig. 1K. A-H were taken with a confocal microscope. Scale bar in A-D is 200 µm. Scale bar in E-H is 10 µm. Each experiment in A-I was performed three times independently except the one where PEG was added which was performed two times. Each data point shown in J comes from an independent experiment.

Next, we developed a minimal model to rationalize the transition from extensile bundles to contractile asters. We took a self-assembly approach where polar and nematic interactions compete to dictate the steady-state structure of the kinesin-microtubule network. Polar interactions depend on the ability of the motor clusters to end-accumulate. If these multivalent clusters form a motor cap on a microtubule, they will naturally recruit other microtubules that will end-accumulate. These end-bound microtubules are free to splay but cannot slide, and hence will form an aster [30,42]. Nematic interactions depend on the ability of the microtubules to stay aligned. In this state, the microtubules are free to slide but cannot splay, and hence will form an extensile bundle.

Using the well-known depletion interaction [63,64], we modeled the effective strength of the alignment interaction between two microtubules as:

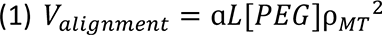

where ɑ is a proportionality constant, - is the average microtubule length, [*PEG*] is the concentration of PEG in the system, and ⍴*_MT_* is the number density of microtubules. The quadratic scaling for ⍴*_MT_* results from the pairwise interaction between microtubules. Similarly, we modeled the effective strength of the end-adhesion interaction as:

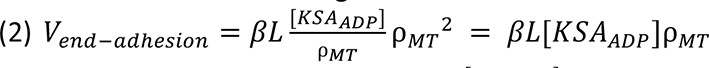

where β is a proportionality constant and 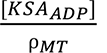 is the number of nonmotile, ADP-bound motor clusters per microtubule. In this model, the number of bound motors increases linearly with microtubule length, and thus the strength of end-adhesion interaction increases linearly with *L*. This scaling is motivated by the experimental observation that the probability of having a motor cap increases with microtubule length, which is reminiscent of an antenna model [32,53] (**Fig. S2G**). As the run length of the motor clusters was larger than the microtubule length (**Fig. S4**), we estimated that all motors walk towards the microtubules’ end and therefore only contribute to the end-adhesion interaction. Further, we considered that only ADP-bound motors that are attached at the end of a microtubule contribute to the end-adhesion. Considering an alternative model where ATP-bound motor clusters contribute to end-adhesion was incompatible with experimental measurements (**Supplementary Materials, section 5, Fig. S10**). We further assumed that the binding of ATP to the motor clusters follows a simple binding equilibrium characterized by a dissociation constant *K_d_*. The concentration of ADP-bound clusters is given by:

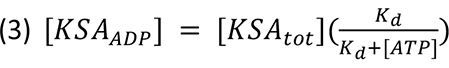

where [*KSA_tot_*] is the total concentration of motor clusters, and *K_d_* is the dissociation constant for ATP binding to the motor clusters (*i.e.* the concentration of ATP where half of the motors have ATP bound).

In this framework, the phase boundary between the bundle and aster phases should lie where the nematic alignment interaction strength (**Eq. 1**) equals the polar end-adhesion interaction strength (**Eq. 2**), providing the following prediction for the phase boundary in the ATP-motor cluster plane:

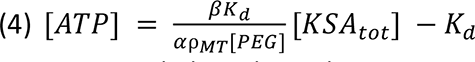

The experimental phase boundary separating the extensile bundle phase from the contractile aster phase in the ATP-motor cluster plane was linear in the range of concentrations explored (**Fig. 1K**), in agreement with the model. Fitting the experimental phase boundary to the model prediction (**Eqn. 4**), provided estimates of the model parameters 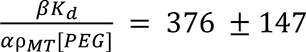 and *K_d_* = 14.5 ± 25 µ*M* (fit values ± 95% confidence intervals).

We independently estimated *K_d_* = 78 ± 41 µM by measuring how the steady state speed of extensile gels varies with ATP concentration (**Fig. S11, Supplementary Materials, section 4c**), which is in agreement with reported values for the same K401 construct, albeit under different experimental conditions *K_d_* = 96.4 ± 25 µ*M* [65]). Further work is needed to understand why the effective *K_d_* derived from fitting the model differed from independent measurements.

Furthermore, the model predicts that the phase boundary in the PEG-motor clusters phase diagram is linear, and the previously measured fit parameters set the boundary’s slope. We overlaid the fit-free prediction for the phase boundary with the experimental data points and found excellent agreement between experiments and theory (**Fig. 4J**). Finally, the model predicts that the slope of the phase boundary in the ATP-motor cluster plane scales inversely with the microtubule density, which was consistent with experimental observations that increasing microtubule number density induced a transition from contractile asters to extensile bundles in PEG-based active gels (**Fig. S3A**).

This self-assembly framework can be modified to account for aligning interactions produced by the microtubule-specific crosslinker PRC1 instead of the depleting agent PEG. We modeled the effective strength of the alignment interaction between two microtubules as:

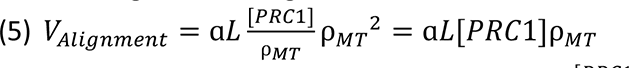

where the number of crosslinkers per microtubule 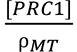 replaces the concentration of PEG as PRC1 needs to bind to a microtubule to induce alignment, while PEG does not. The theory predicts that the new phase boundary in the ATP-motor cluster plane is now independent of the density of microtubules ⍴*_MT_*:

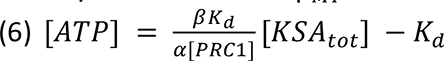

We assembled PRC1-based active microtubule gels in a contractile phase and systematically increased the number density of microtubules from 0.33 mg/mL to 10 mg/mL (**Fig. S3B-C)**. We found that the emergent dynamics were always contractile in that range of concentrations, consistent with Eq. 6. Therefore, while both PRC1 and PEG induce nematic alignment of microtubules, this minimal model accounts for the different ways they interact with microtubules, explaining their distinctive impact on the self-assembly of microtubule-based active gels.

Despite the initial complexity of the five-dimensional organization phase space – ATP, motor clusters and PEG concentrations, microtubule number density, and microtubule length - this model provides a simple design rule to assemble active microtubule gels in targeted configurations. In this framework, a single control parameter given by the ratio of *K_d_* and the various component concentrations: 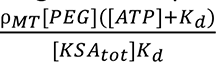 dictates the self-assembly of the emergent structures in PEG-based active gels. Estimating ⍴*_MT_* from the known tubulin concentration, the number of protofilaments in a microtubule, and the measured length distribution, we rescaled all the experimental data points from ∼ 300 independent experiments (**Fig. S12**) onto a single 1D phase diagram. We computed the probability of being in a globally contractile, aster, or extensile phase as the ratio 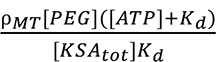 increased (**Fig. 5B**). The model predicts that the phase boundary between the extensile and aster phase is set by the critical number density 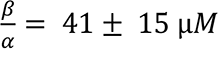 (95% confidence interval on linear fit). Below this critical density, our model predicts that microtubules will form asters, while above they will form bundles. We find that the number density *c*^∗^ where extensile and contractile phases are equiprobable lies within the 95% confidence interval of the fitted value for 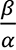(*c*^∗^ = 41 µ*M*, **Fig. 5B**). Excellent agreement between theory and experiments suggests that a self-assembly process governed by the competition between polar and nematic alignment interactions describes the extensile-to-contractile transition (**Fig. 5A**).

**Figure 5:**
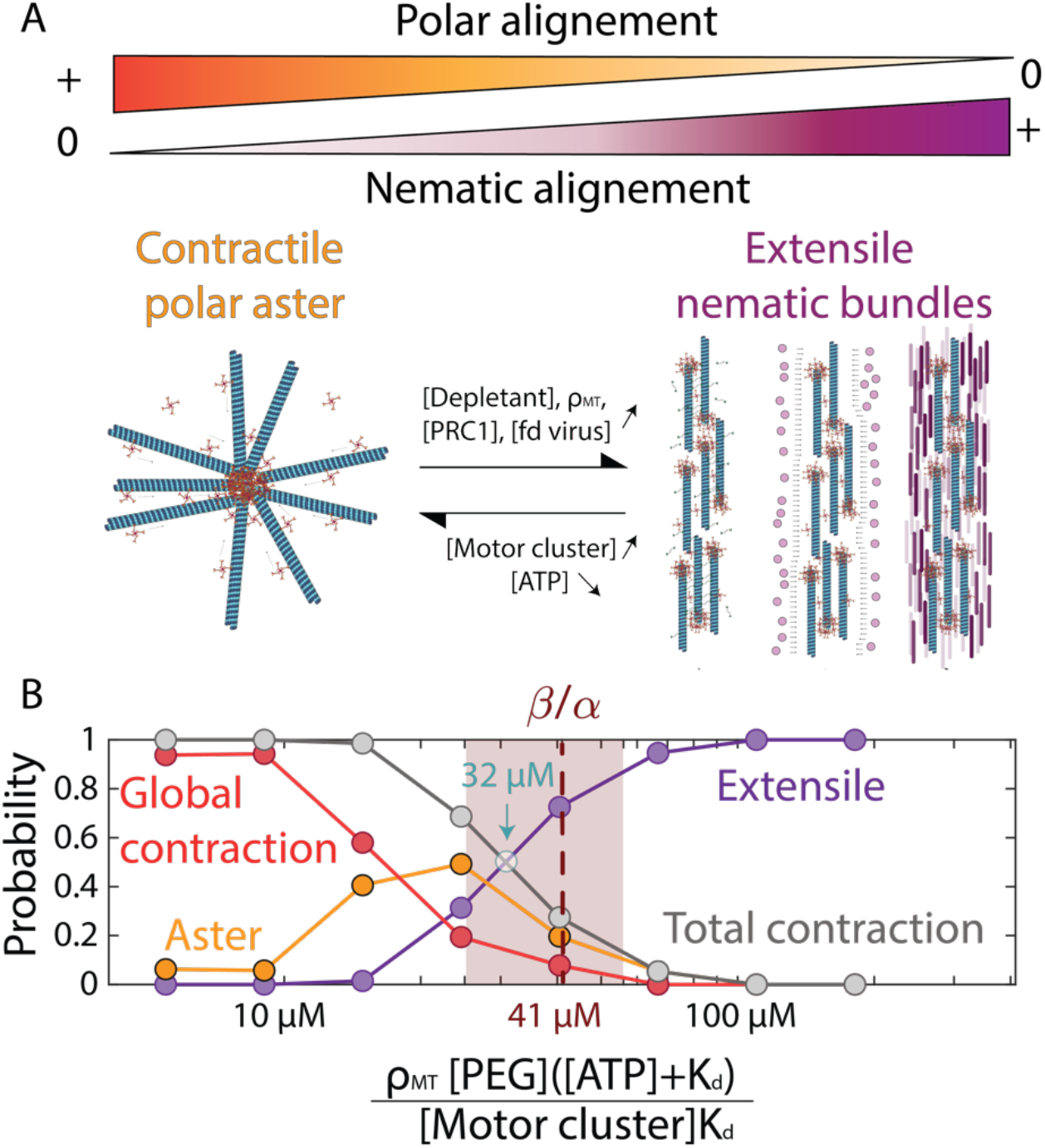
Nematic alignment competes with polar sorting to control the self-assembly into extensile bundles or contractile asters. A) Schematic representation of a contractile aster composed of polarity-sorted microtubules. Increasing the strength of nematic alignment by increasing the concentration of depletant, microtubule, PRC1 crosslinker, or *fd* virus can trigger a structural transition from contractile aster to extensile bundles. Decreasing the concentration of motor cluster or increasing the concentration of ATP decreases the probability of cap formation, which decreases the strength of polar end-adhesion interactions. At the macroscale, this leads to a transition from contractile asters to extensile bundles. B) Relative probability of finding the active gel in the extensile, aster, or globally contractile phases as the rescaled concentration 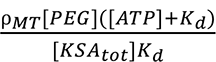 increases. Disks show experimental data points. The raw 2D phase diagrams are shown in Fig. 1K, 4J, and **S12**. Above a critical microtubule density 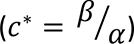, the active gel evolves on average from a contractile to extensile phase. The experimental estimate of *c*^∗^ where extensile and contractile phases are equiprobable is 32µM. The dashed line shows the fitted value for *c*^∗^ = 41 µ*M* and the brown area shows the corresponding interval of 95% confidence for *c*^∗^ [25-56µM]. Both are estimated by fitting the phase diagram in Fig. 1K. (In B, N=311 independent experiments, for [ATP]=20-150µM, [Motor cluster]=40-500nM, [PEG]=0.2-0.6mM, i.e 0.4-1.2 %/% vol, [MT]=2.7-16nM).

## Discussion

To summarize, we bridged the length scale gap between the molecular interactions and the emergent macroscopic structure, explaining the transition from extensile bundles to contractile asters in active gels composed of stabilized microtubules, multivalent clusters of kinesin-1 motors, and either a depletant or microtubule-specific crosslinkers. We identified kinetics of motor-microtubule interactions that lead to either uniform motor distribution along microtubules or the end-accumulation of motors, which relate in part to the emergence of extensile bundles or contractile asters. We show that the dynamics of end-accumulation at the single filament scale sets the macroscopic emergent timescale of aster formation. Further, we show that end-clustering is insufficient to explain fully the extensile-to-contractile transition. Nematic interactions driven by depletion, crosslinking, or excluded volume interactions can outcompete polar sorting to favor extensile bundles over polarity-sorted contractile asters, even in conditions where motor clusters end-accumulate. A minimal self-assembly model based on the competition between microtubule end-clustering driven by end-accumulated motors and nematic alignment captures the experimental trends. We further verified predictions of the model that differentiated between depletant and crosslinker-based active gels and demonstrated that a single control parameter given by the ratio of the concentrations of the components sets the structural transition from extensile bundles to contractile asters.

We consider these findings in the context of previously published results on the extensile-to-contractile transition in cytoskeletal active gels. First, while previous work on kinesin-microtubule gels focused on the role of side binding versus end-binding motors [43], direct observation of both the bulk organization and motor cluster distribution on single microtubules reveals that nematic bundles can arise even when motors efficiently end-accumulate.

Second, several mechanisms have been proposed to account for the extensile or contractile nature of biomimetic active matter. Recent theoretical work proposed that nonlinear elasticity can rectify the activity of an extensile dipole, leading to bulk contraction [66-68]. For actomyosin gels, non-linearities in actin filament mechanics have been put forward as one mechanism leading to contraction. While similar emergent mechanical properties have been suggested in visco-elastic gels composed of kinesin-microtubules bundles dispersed in a passive actin network [18], our multiscale approach favors that the asymmetric distribution of motors leads to aster formation in kinesin-microtubule-based active gels. Our work highlights that protein-protein interaction kinetics plays a crucial role in setting the macroscale emergent dynamics.

Third, our work investigates how the contractile-to-extensile transition is impacted by crowding, microtubule density, and microtubule length, all of which were recently explored in computer simulations of kinesin-microtubule networks [69]. While we agree that crowding promotes extensile bundles over contractile asters, we report contradicting results regarding the impact of microtubule length and their number density: increasing microtubule density leads to extensile bundles in PEG-based networks and increasing microtubule length does not impact the transition. The reason for the latter is that longer microtubules favor nematic alignment and end-accumulation equally, but only when their length is shorter than the motor clusters’ run length. When microtubules are longer than the run length, our model predicts that increasing microtubule length only enhances the alignment interaction, leading to a transition from contractile asters to extensile bundles, thus corroborating the results of the computer simulations [69]. Our results are also consistent with how microtubule networks powered by end-accumulating kinesin-4 motors self-organize in the presence of a depletant [34].

Finally, while depletant has been a long-time favorite bundler for microtubule-based active gel [4], PRC1 crosslinkers have recently been used to assemble extensile active gels [6,70,71] and active composite materials [18,62]. Our work highlights for the first time the similarities and differences of how depletants and crosslinkers impact the self-organization of microtubule-based active gels.

This work illustrates how to leverage simple concepts from self-assembly to understand complex emergent structures and dynamics in an out-of-equilibrium material powered by chemical reactions. One important takeaway of the theory is that asters nucleation and polarity sorting are not the consequence but rather the cause of local contractility. However, it is unclear why this minimal model captures the extensile-to-contractile transition so well (**Fig. 5B**). Indeed, this mean-field theory only considers pairwise interactions and lacks any higher-order cooperativity between microtubules, or between microtubules and motor clusters [33,72,73]. Second, this theory has no intrinsic timescales, hence no dependence on reaction rates which are crucial for explaining the interactions between kinesin and microtubules. Third, this model cannot describe the scale-dependent architecture of the active gels. When end-adhesion dominates, microtubules with motor caps are predicted to form micelle-like structures with a radius of one microtubule length [34]. However, the aster sizes observed here are orders of magnitude larger than the average microtubule length, which clearly shows that the model cannot give information about the hierarchical organization that leads to the observed mesoscale structures (**Fig. 1I-J**). Consequently, this model does not describe the percolation transition from asters to globally contractile gels [74]. Finally, we expect that the critical ratio delineating the extensile and contractile phases will be different for different motors, cluster valency, or buffer conditions (**Fig. 5B**). One piece of evidence is that adding salt eliminates any contraction without changing protein composition (**Fig. S5**).

These results further highlight that motor clusters’ valency and processivity dictate in part the self-organization of the active gel at the macroscale. Processivity is required to drive end-accumulation [32] and aster formation. No contraction had been observed with light-activated K365 dimers as they are non-processive by design (**Fig. S6**). In contrast, the other constructs potentially form asters because the motor clusters are multivalent, hence highly processive [40], and readily able to end-accumulate (**Fig. S13)**. These results underscore the importance of paying close attention to the design of the motor cluster, as they significantly impact the self-organization of the cytoskeletal active gels.

Taken together, this work offers a blueprint for the predictive assembly of cytoskeletal active matter, particularly for building multifunctional active materials that can be programmed into distinct or coexisting far-from-equilibrium structures.

## Acknowledgment

We thank Shibani Dalal, director of the Brandeis Biomaterial Facility, for help with protein purification. B.N. thanks Saptorshi Ghosh for the discussion on data analysis. We thank Daichi Hayakawa for providing 6 helix bundle DNA origami rods and Salman Alam for providing the *fd* viruses. We thank Linea Lemma for the opto-K365 constructs and Bennett Sessa for comments on the manuscript. B.N. thanks Ian Murphy for help in native gels.

B.N. and G.D. acknowledge support from the NSF CAREER award DMR-2047119. P.J.F. and A.B. acknowledge support from the Brandeis NSF Materials Research Science and Engineering Center (MRSEC) DMR-2011846. We also acknowledge the use of the optical, microfluidics, and biomaterial facilities supported by NSF MRSEC DMR-2011846. A.B. acknowledges support from NSF-DMR-2202353.

## Methods and Materials

### 1. Protein purification protocols

#### Microtubules

Tubulin dimers were purified from bovine brains through two cycles of polymerization– depolymerization in high molarity PIPES (1,4-piperazindiethanesulfonic acid) buffer [1]. Fluorophore-labeled tubulin was prepared by labeling the purified tubulin with Alexa-Fluor 647-NHS (Invitrogen, A-20006) [2]. GMPCPP (Guanosine 5’-(α,β, methylenetriphosphate)), a non-hydrolyzable analog of GTP, was used to stabilize the dynamic instability in the MTs. The polymerization mixture consisted of 80 µM tubulin (with 3% fluorescently labeled tubulin), 0.6 mM GMPCPP, and 1 mM dithiothreitol (DTT) in M2B buffer (80 mM PIPES, 1 mM EGTA, 2 mM MgCl2). After adding all the components, the mixture was incubated at 37 °C for 30 min, and subsequently for 6 h at room temperature (25 °C). The stock concentration was 8 mg/mL of tubulin. MTs were aliquoted in small volumes (5 µL), flash-frozen in liquid nitrogen, and stored at −80 °C.

#### Crosslinkers—PRC1-NSΔC

The PRC1-NSΔC (MW: 58 kDa) is a truncated version of the microtubule crosslinking protein PRC1. The deletion of the unstructured C-terminus of the PRC1 protein results in reduced interactions between adjacent crosslinkers. PRC1-NSΔC plasmids were a gift from Radhika Subramanian [3]. The protein was expressed and purified from Rosetta BL21(DE3) cells via ÄKTA pure FPLC system using an established protocol described in [3]. These proteins were flash-frozen with 40% sucrose in 50 mM Potassium phosphate (pH 7.0), 500 mM Potassium chloride, 5% glycerol, and 3 mM DTT and stored at −80 °C. The final concentration of PRC1 was measured by a Bradford assay.

#### K401 Molecular motors

K401-BIO-6xHIS (processive motor, dimeric MW: 110 kDa) are biotinylated kinesin constructs derived from N-terminal domain of Drosophila melanogaster kinesin-1, truncated at residue 401 and labeled with six histidine tags for Nickle affinity purification. K401-BIO-6xHIS plasmids were a gift from Zvonimir Dogic [2]. The motor proteins were transformed and expressed in Rosetta (DE3) pLysS cells and purified following established protocols described in [4]. Briefly, clarified lysate obtained from the cells was bound to a HiTrap Chelating HP column (GE Healthcare) in low imidazole buffer (50 mM PIPES, 20 mM imidazole, 4 mM MgCl2, 50 μM ATP, 2 mM DTT, pH 7.2). Protein was subsequently eluted by an imidazole gradient that ranged from 20 mM to 500 mM. The protein was stored at 1mg/mL in 50 mM PIPES, 20 mM imidazole (pH 7.2), 4 mM MgCl2, 2 mM DTT, 50 μM ATP and 36% sucrose buffer at −80 °C. The final concentration of kinesin was measured by Bradford assay.

We used tetrameric streptavidin (ThermoFisher, 21122, MW: 52.8 kDa) to assemble clusters of biotinylated kinesins (KSA). To make K401-streptavidin clusters, 5.7 µL of 6.6 µM streptavidin was mixed with 5 µL of 6.4 µM K401 and 0.5 µL of 5 mM DTT in M2B. This mixture was incubated on ice for 30 minutes.

#### Light-activable motors: K365-iLID and K365-micro

K365-iLID and K365-micro were designed by Linnea Lemma and Tyler Ross and were purified following the protocol described in ref. [5,6]. Briefly, two chimeras of the D. melanogaster kinesin K365, namely, K365-iLID and K365-micro, were expressed in Escherichia coli Rosetta 2(DE3)pLysS cells and purified using ÄKTA pure FPLC system. MBP domain was cleaved with TEV protease (Sigma Aldrich). These proteins were flash-frozen with 40% glycerol in a solution containing 50 mM Sodium Phosphate (pH 7.2), 20 mM imidazole, 4 mM MgCl_2_, 250 mM NaCl, 5 mM BME, and 50 μM ATP. The proteins were stored at a concentration of 1mg/mL at −80 °C.

#### Preparation of DNA origami rods

DNA origami rods were prepared via established protocols [7]. Briefly, each DNA origami particle is folded in a solution consisting of 50 nM of p8064 scaffold DNA from Tilibit, 200 nM each of staple strands obtained from IDT, and 1xFoB5 (5 mM Tris Base, 1 mM EDTA, 5 mM NaCl, and 5mM MgCl2). After incubation at 65◦C for 15 minutes, the folding mixture is annealed from 54◦C to 51◦C at 1 hour /◦C, using a Tetrad thermocycler from Bio-Rad. To assess the success of folding, the mixture is separated using agarose gel electrophoresis at 110 V for 1.5 to 2 hours in a cold room maintained at 4◦C. The resulting gel is scanned at 100 µm resolution using a Typhoon FLA 9500 laser scanner from GE Healthcare. Following folding, DNA origami particles are purified to eliminate excess staples through PEG purification. Folded solution, 1xFoB35 (5 mM Tris Base, 1 mM EDTA, 5 mM NaCl, and 35mM MgCl2), and PEG buffer (5 mM Tris Base, 1 mM EDTA, 15% PEG8000, 500mM NaCl) is mixed with a ratio of 1:1:2. The mixture is centrifuged for 90 minutes at 4.6krcf. After removing the supernatant, the pellet is carefully washed in 1xFoB5 twice, before resuspending in proper volumes of 1xFoB5.

#### Fd virus purification

Standard biological procedures were followed to grow and purified filamentous fd virus [8,9], using Xl1-Blue as the host E. coli strain. Virus samples were stored in M2B buffer. Concentration of the virus was determined using a Nanodrop.

### 2. Assembling 3D active gels for bulk experiments

The active gels are composed of:

- Alexa 647-labeled GMPCPP stabilized MTs with an exponential distribution of lengths with an average of 1.5 µm, unless otherwise specified,
- Multi-motor kinesin complexes self-assembled from tetrameric streptavidin and two-headed biotinylated kinesin (K401-Bio) or single-headed biotinylated kinesin (K365-Bio)
- MT bundling, (0.6-2)% (w/v) PEG (polyethylene glycol, 20 kDa, Sigma) or 100nM PRC1, that passively crosslink antiparallel MTs, but still allow interfilament sliding,
- ATP (Sigma)
- An ATP regeneration system: phosphoenol pyruvate (53 mM PEP, Beantown Chemical, 129745) and pyruvate kinase/lactate dehydrogenase enzymes (2.8% v/v PK/LDH, Sigma, P-0294). For experiments with PRC1, we used 26mM of PEP.
- An oxygen scavenging system comprised of glucose (18.7 mM), DTT (5.5 mM), glucose oxidase (1.4 µM), and catalase (0.17 µM) was used to decrease photobleaching.

For consistency, a large volume of premix was prepared, aliquoted, and snap-frozen in liquid nitrogen for each phase diagram (Fig. 1, 4, S12). Frozen MTs (stored at −80 °C) were thawed immediately before use in an experiment. All the experiments were performed at room temperature. Active networks composed of MTs (1.3 mg/mL), ATP (1420 µM), and K401 clusters (120 nM) remain active for 6–8 h.

#### 2.a Flow chamber assembly for bulk experiments

All flow chambers have the same dimensions (H = 80 µm, W = 3 mm) and were assembled using two glass slides spaced by a layer of parafilm. The glass surfaces were coated with an acrylamide brush to resist non-specific protein adsorption [2]. Parafilm spacers were cut and placed between the two glass surfaces followed by a mild heat treatment at 60 °C to melt parafilm so it can bind to the glass surfaces. The active mixture was loaded into each channel type by capillarity and sealed with an UV-curing optical adhesive (NOA 81, Norland Products Inc.).

#### 2.b Flow chambers for single microtubule studies

The flow chambers were constructed as previously described [10]. In brief, we built a chamber by assembling two silanized coverslips (VWR, Germany) on the glass spacers cut out of a coverslip (thickness ≈ 0.140 mm). The chamber was incubated with 16% (v/v) glutaraldehyde (Electron Microscopy Sciences) for 30 minutes. For single molecule studies, the MTs were polymerized from a solution of 80 µM of 20% Alexa647-labeled tubulin. Microtubules (115 nM tubulin) were incubated in the flow chamber at room temperature. The remaining exposed surface was blocked for 5 minutes with 6% Pluronic F-127 (w/v) (Sigma P2443. MW: 12.5 kDa) and 15mg/ml of Bovine Serum Albumin (BSA, Sigma) solution.

The imaging buffer consisted of M2B supplemented with 4 mg/ml BSA, (0-2)% (w/v) PEG (polyethylene glycol, 20 kDa), 3mM MgCl2, ATP (ranging from 5 to 200 µM), ATP regeneration system, 2mM Trolox, an oxygen scavenging system, and motor clusters ranging from 40 to 800 nM (Alexa 568 or Alexa 488 labeled Streptavidin).

#### 2c. Native Page gels protocol

We used the blue native-PAGE method to confirm that kinesin-1 K365 ilid and kinesin-1 K365 micro constructs do not spontaneously dimerize in dark.

Blue native-PAGE was performed by adding proteins to 4x RunBlue Native Sample Buffer (expedeon). Samples were subsequently loaded onto NuPAGE™ 3–8% Tris-Acetate (Invitrogen) and NativePAGE™ 4–16%, Bis-Tris gradient gels (Invitrogen) and run at 150 V in Tris-glycine and TBE buffers, respectively. NativeMark™ Unstained Protein Standard (Invitrogen) were loaded to predict the size of detected protein species. The experiments were performed at 4oC in dark.

The gels were subsequently stained in a solution consisting of 30% methanol, 10% glacial acetic acid, and 0.02% Coomassie R-250, then placed on a shaker for 30 minutes. Afterward, the gels were destained twice for six hours each time in a solution of 20% methanol and 10% glacial acetic acid on a shaker before scanning.

### 3. Microscopy

#### Widefield microscopy

The Alexa 647-labeled MT networks were imaged using an inverted widefield microscope (Nikon Ti-E or Ti2) with a fluorescent filter (Semrock Cy5-4040C), a SOLA light engine (Lumencor), 10x (Nikon Pan Fluor, NA 0.3) or 4x objective (Nikon Pan Fluor, NA 0.13) and a CCD or a sCMOS camera (Andor Clara E, Hamamatsu orca flash 4.0). The illumination and the data acquisition were controlled by micro-manager (µManager, Version 2.0.0-gamma [11]). All the measurements were performed at room temperature.

#### Confocal microscopy

Imaging of motor cluster transport and accumulation on microtubule plus ends was performed using Leica Application Suite X (LAS X) control on a laser scanning confocal microscope (TCS-SP8, Leica Microsystems GmbH) equipped with photomultiplier tubes.

For excitation, we used a 638 nm laser diode for Alexa Fluor 647 labeled microtubules, and 488 nm or 552 nm laser diodes for Alexa Fluor 488 or Alexa Fluor 568-labeled motor clusters, respectively. Images were acquired using an oil-immersion 63x objective (HCX PL Fluotar, NA =1.4).

### 4. Image and Data Analysis

#### 4.a Single filament cap analyses

The probability distribution of motor caps at microtubule ends was measured using a custom-written MATLAB program. We collected timelapse series of dual-channel confocal images where both tubulin and motor clusters were labeled. We first segmented microtubules after excluding microtubules at the edges of the image and overlapping microtubules. The motor channel was scanned along the length of each segmented microtubule for the presence of a motor cap by comparing the maximum intensity to the average intensity of the background. If that ratio was greater than 2.5, then a cap was detected. The probability distribution of caps is calculated by taking the ratio of the number of caps to the number of microtubules for each image.

#### 4.b How to determine if the system is in the extensile phase, aster phase or the globally contractile phase?

Time series of samples with fluorescent microtubules were inspected by eye. Asters are identified as local and sustained densification of the microtubules. In the absence of contraction over the entire lifetime of the sample (10-12h), the dynamics is labeled as extensile. If any asters form during the lifetime of the sample, the gel is labeled as an aster phase.

#### 4.c Determination of K_d_

We measured !_!_ by quantifying the ATP dependence of active flow speeds. We measured the average speed of active flows as a function of ATP over three decades of ATP concentration (Fig.S11).

Premixes of extensile gels were prepared with different ATP concentrations between 20 and 1420 µM using 110 nM Motor clusters and 1.33mg/mL of tubulin. For each ATP concentration, the experiment was repeated four independent times.

Flow fields of microtubule bundles were measured using MATLAB-based open-source particle image velocimetry (PIV) [12]. An interrogation window size of 64 pixels = 82.56µm was selected with 50% overlap.

#### 4.d Estimation of the microtubules number density

For a microtubule of 1.5μm length, there are about 13 protofilaments, hence 2440 tubulin dimers. If one tubulin dimer weighs 100 kDa then one microtubule weighs 244,000 kDa. Therefore, for an active network containing 1.33mg/mL of tubulin, the microtubule number density is 5.45nM (if one considers that all the tubulin will polymerize into microtubules).

#### 4.e Run length measurement

We measured the run length, i.e. the distance motors travel along microtubules before detaching or stopping. We first segmented the microtubules, then analyzed the tracks of the kinesin clusters using kymographs. We report the run lengths of 216 events in **Fig. S4C.**

#### 4.f Measurement of microtubule length distribution

To characterize the length distribution of the microtubules, the MTs were diluted to 3000X with antioxidants, Trolox, and a 2.5% (w/v) solution of dextran (MW 500 kDa). The dilution helps to prevent overlaps of MTs that complicate automated filament recognition. For imaging fluorescently labelled microtubules, 5µL of the solution was placed between a coverslip and a coverslide. Imaging was performed on a standard fluorescence microscope (Nikon Eclipse Ti microscope) using a high numerical aperture oil objective (Nikon Plan Flour 100X/1.30). We wrote a custom Matlab script that quantifies the length distribution of MTs. Binary images of each MT were fitted with an ellipse. The long axis of the ellipse corresponds to the MT’s length.

### 5. Alternative theoretical model

We present in this section how the extensile/contractile phase boundary would look like if we considered only the ATP-bound motor clusters instead of ADP-bound motor clusters.

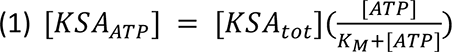

We ended up with the following equation for the phase boundary:

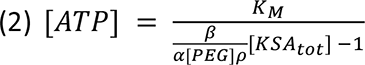

This phase boundary does not agree at all with the experimental data (**Fig. S10**), suggesting that we should consider the ADP-bound motors as contributing to the end-clustering, not the ATP-bound motors.

### 6. Supplementary figures

**Figure S1:**
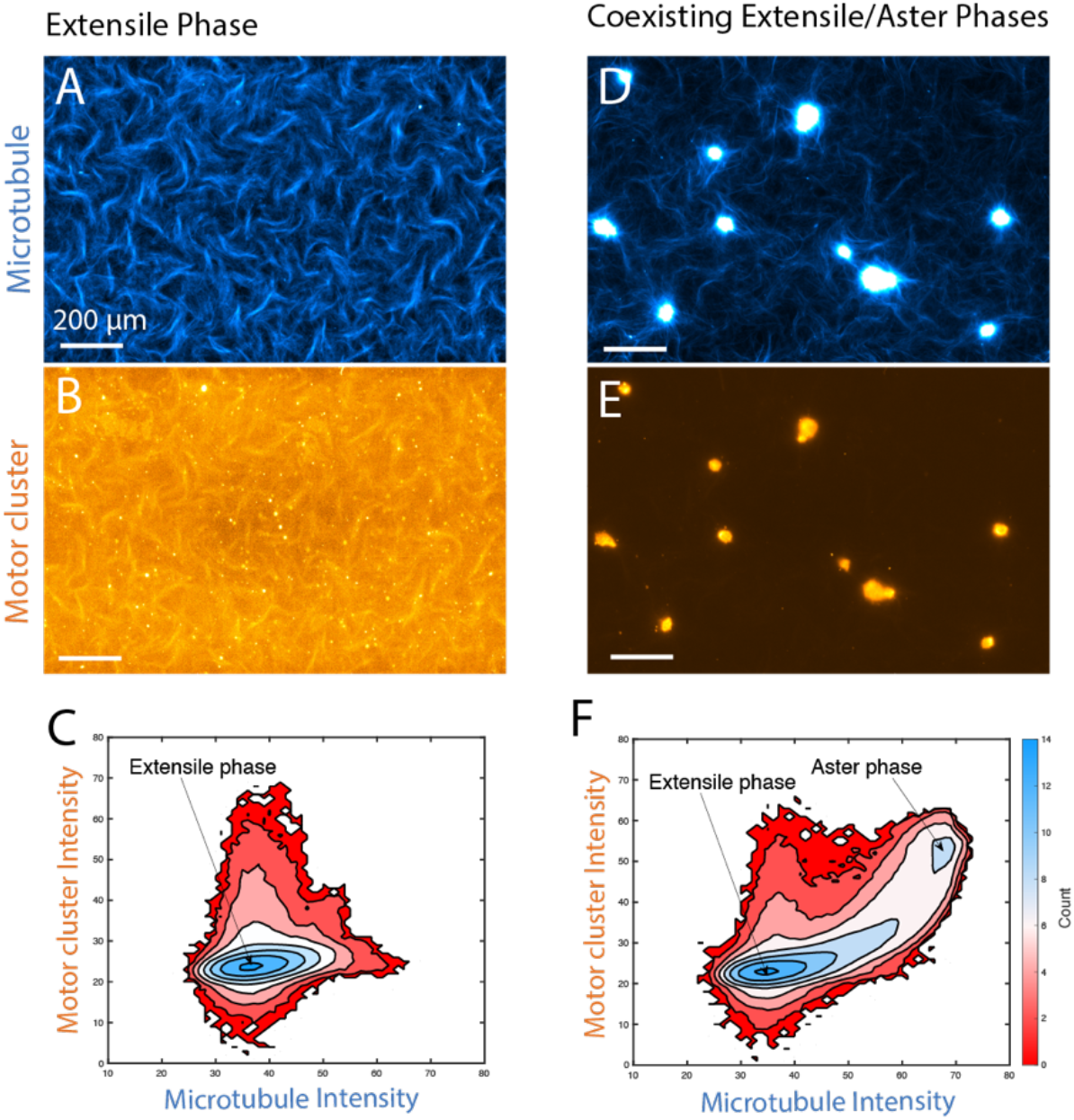
Comparing kinesin cluster to tubulin ratio in the extensile phase and in the aster phase. A) and B) respectively show the tubulin and the motor cluster channels for an extensile gel. C) Probability distribution for the intensity of motor and tubulin in the extensile phase. The ratio of the intensity is constant. D) and E) show the tubulin and the motor cluster channels for asters coexisting in an extensile background. F) Probability distribution for the intensity of motor and tubulin in the coexisting aster/extensile phase. The asters are enriched in motor clusters compare to the bundles. Scale bars in A-B and D-E are 200 µm. C and F have the same color code.

**Figure S2:**
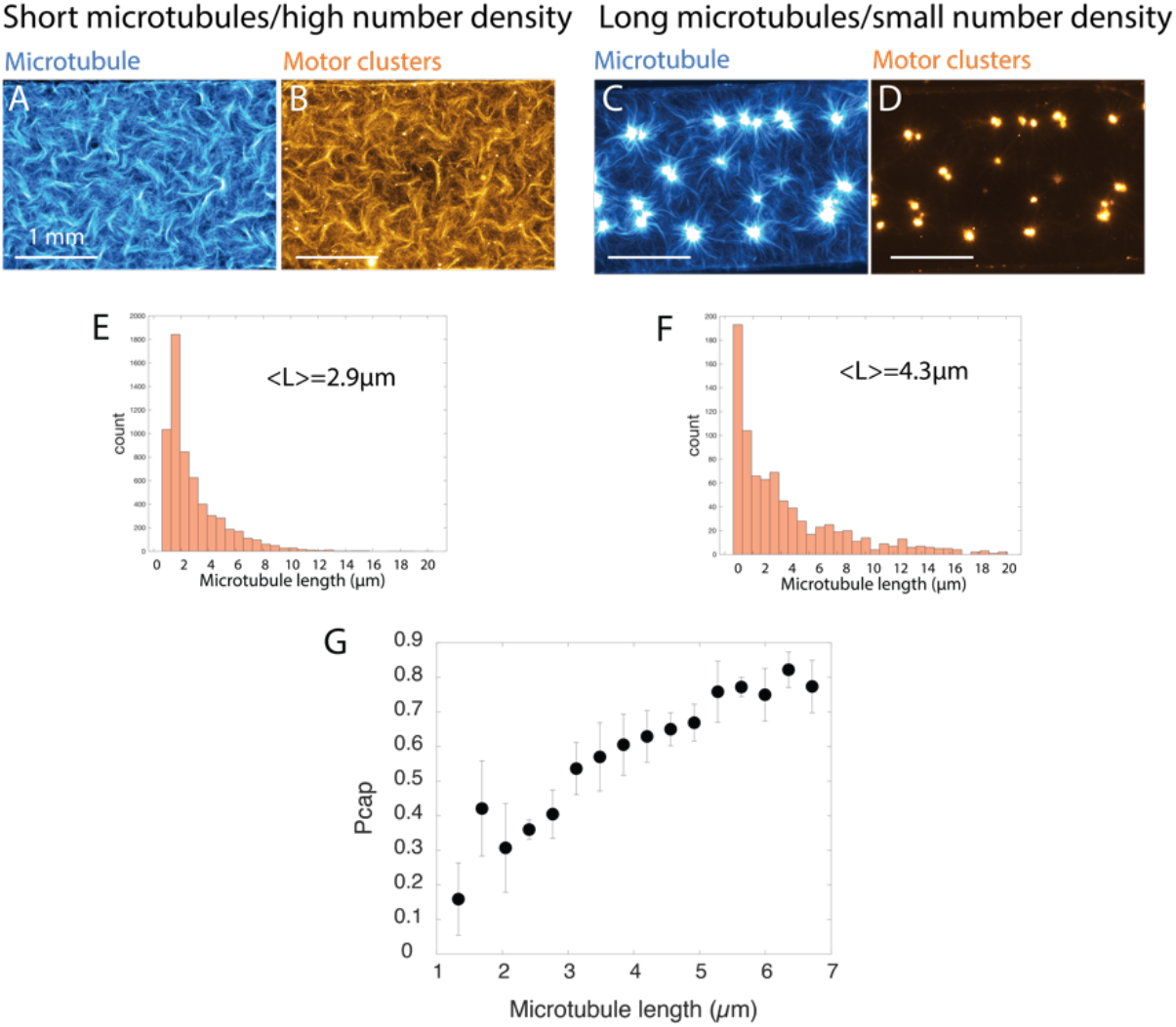
Increasing the microtubule’s average length while keeping tubulin concentration constant leads to a transition from extensile to contractile dynamics in PEG-based active gels. Fluorescent images of tubulin and motor clusters for an average microtubule length of A-B) 2.9µmm and C-D) 4.3µm and a tubulin concentration of 1.33mg/mL. E-F) Probability distribution function for the microtubule length respectively used in A and C. G) Fraction of microtubule with a motor cap as a function of the microtubule length. Longer microtubules have a higher probability of forming a motor cap. [Motor cluster]=110nM, [ATP]=30µM, [tubulin]=1.33mg/mL, [PEG-20kDa]=0.6% (vol/vol). Error bars show standard deviation over binned microtubule of length mean+/-0.5µm. Data shown in G comes from 18 experimental conditions where ATP and motor cluster concentrations were varied over [ATP]=5-150µM and [Motor cluster]=50-800nM. Each experimental condition was repeated 3 times.

**Figure S3:**
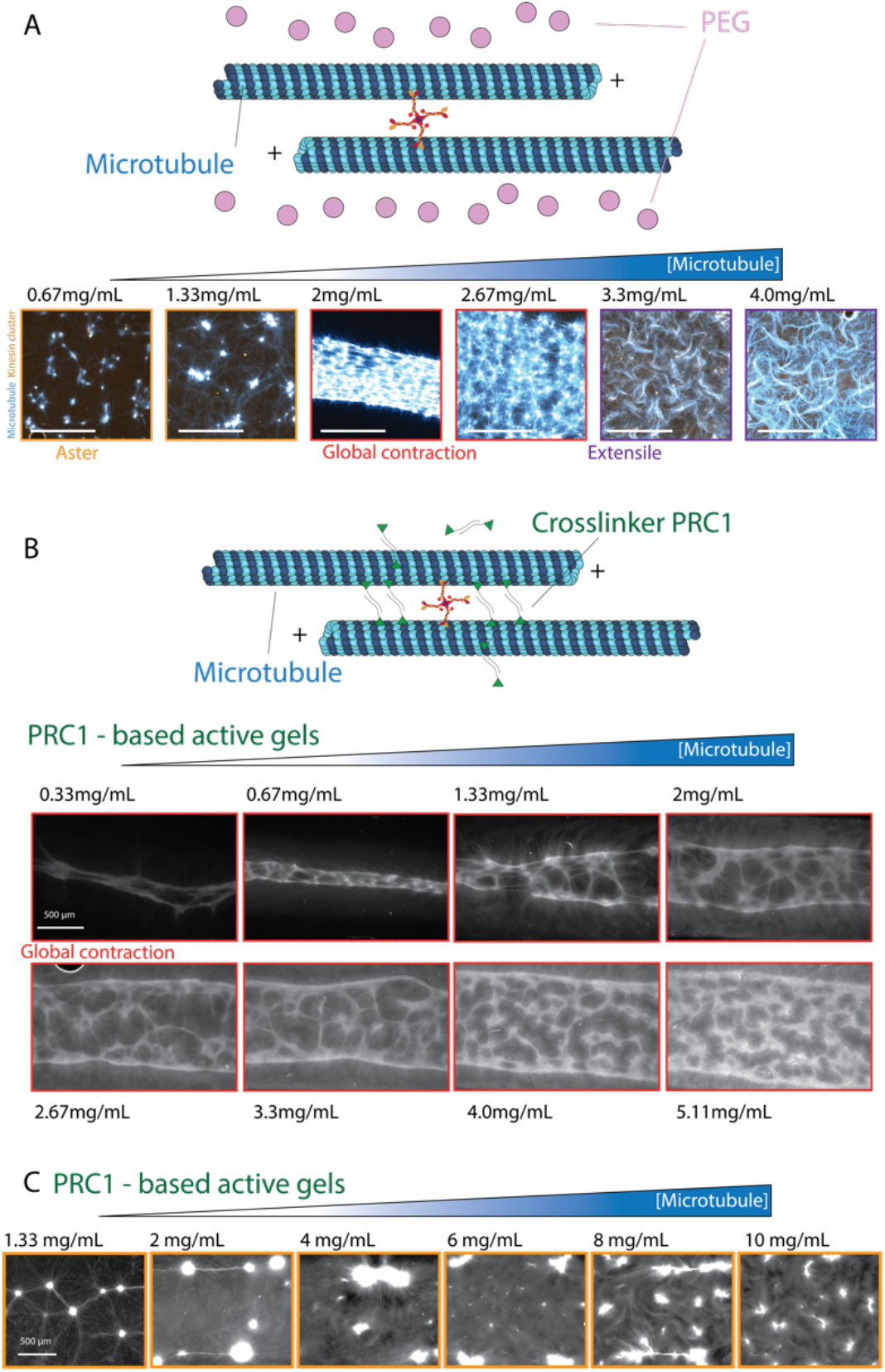
Increasing microtubule concentration leads to a contractile-to-extensile transition for PEG-based active gel and does not impact the self-organization of PRC1-based active gels. A) schematic of a PEG-based bundle and fluorescent images of tubulin (cyan) and motor cluster (orange) when tubulin concentration is increased while keeping the average microtubule length constant. Scale bar is 1 mm, Images were taken with a confocal microscope. [ATP]=30µM, [Motor cluster]=200nM, [PEG-20kDa]=0.6% (vol/vol). B) schematic of a PRC1-based bundle and fluorescent images of microtubules when tubulin concentration is increased from a globally contractile phase. C) Fluorescent images of microtubules when tubulin concentration is increased from an aster phase. In B and C, [ATP]=30µM, [Motor cluster]=380nM, [PRC1]=100nM.

**Figure S4:**
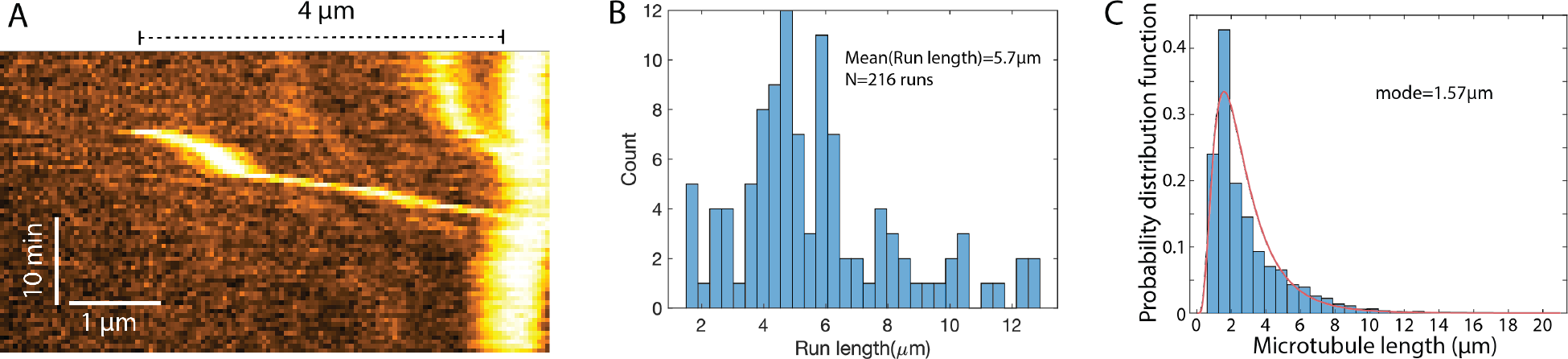
High processivity of clusters of K401 kinesin-1. A) Kymograph (space-time plot) showing a labeled motor cluster binding and moving onto a microtubule. The run length is longer than 4µm. The motor cluster reaches the dense motor cap at the end of the microtubule. B) Probability distribution function of run length of the clusters. C) Probability distribution function of microtubule length.

**Figure S5:**
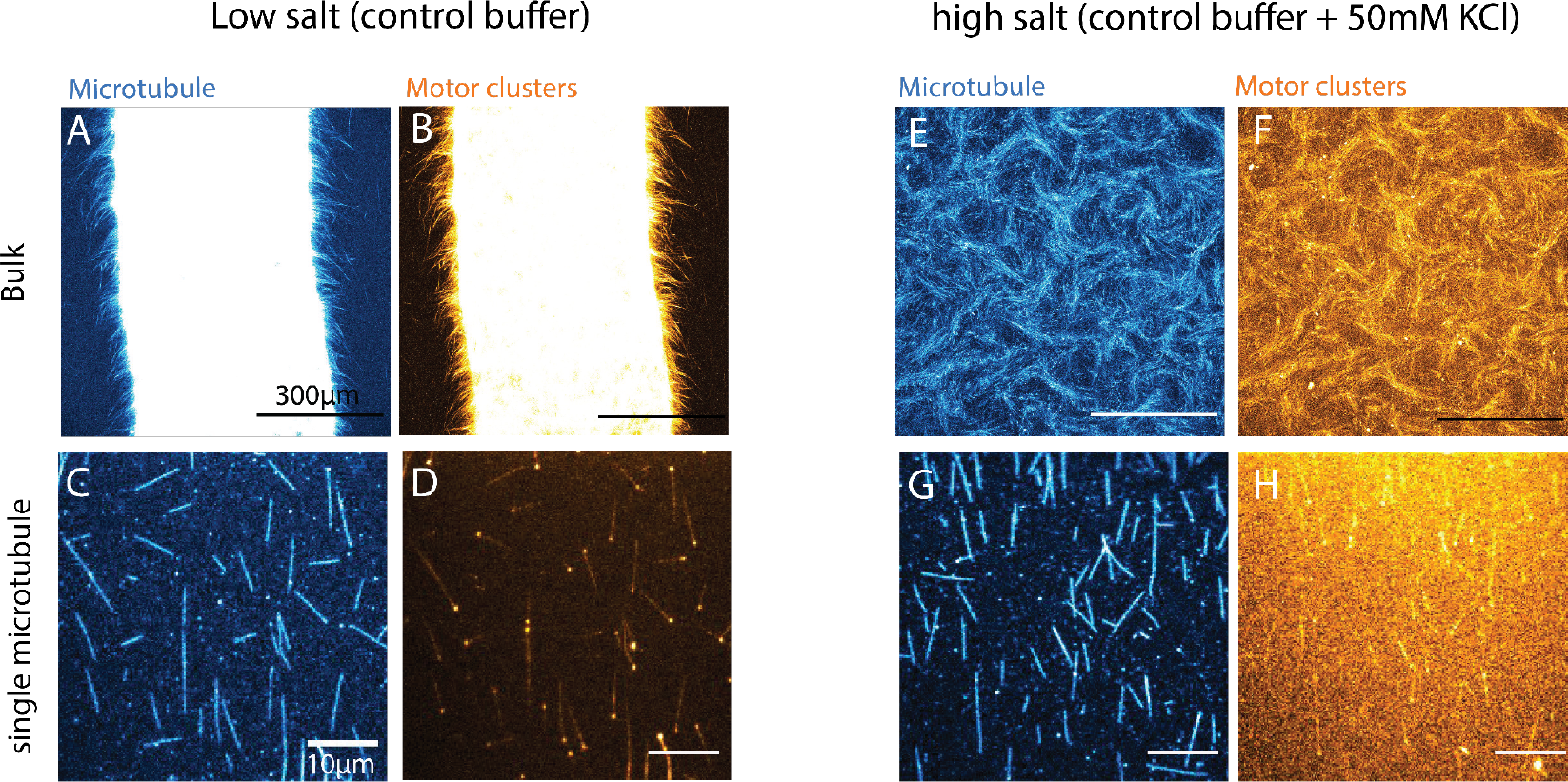
Decreasing the residence time of motors onto microtubules drives the transition from contractile to extensile dynamics. In the absence of KCl (control), the active gels (A) tubulin channel, B) motor channel) globally contracts and motor clusters accumulate at the end of the microtubule (C) tubulin channel, D) motor channel). Adding 50nM of KCl drives bulk extensile dynamics (E) tubulin channel, F) motor channel) and motor caps disappear (G) tubulin channel, H) motor channel). A-H were taken with a confocal microscope. Scale bar in A, B, and F is 300 µm. Scale bar in C, D, G, and H is 10 µm. In all experiments, [ATP]= 30µM, [motor clusters]= 480 nM, [PEG]=0.6% vol/vol. Both bulk and single microtubule experiments were performed in 3 independent replicates.

**Figure S6:**
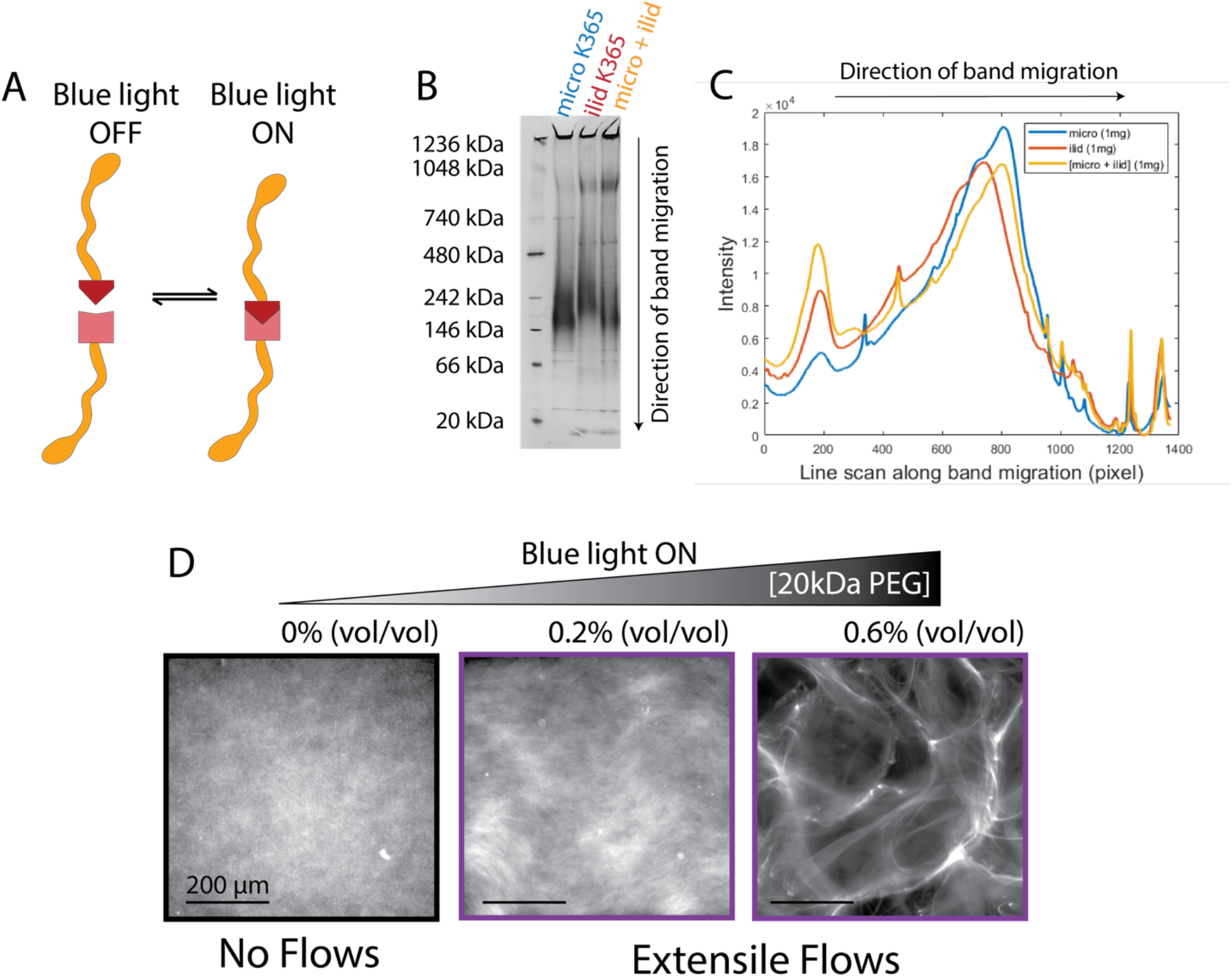
Dimers of K365 do not form asters. A) schematic of the light-dimerizable single-headed opto-K365 motors. When the light is off, the motors are not dimerized. When the blue light is on, the motors dimerize. B) Native gel experiments demonstrate that the motors do not dimerize in the absence of light. Native gel were repeated 4 times on independent samples. C) Quantitative analysis of the native gel showing that the motors do not dimerize or aggregate in the absence of light. D) Fluorescent pictures of the active gels composed of 500nM of each iLid and micro motors. In the absence of PEG, no motion is observed. Adding PEG increases the depletion interaction and extensile motion is detected. ([ATP]=30µM, [MT]=1.33mg/mL, [microK365]=500nM, [IlidK365]=500nM). Experiments in Dwere performed in 3 independent replicates.

**Figure S7:**
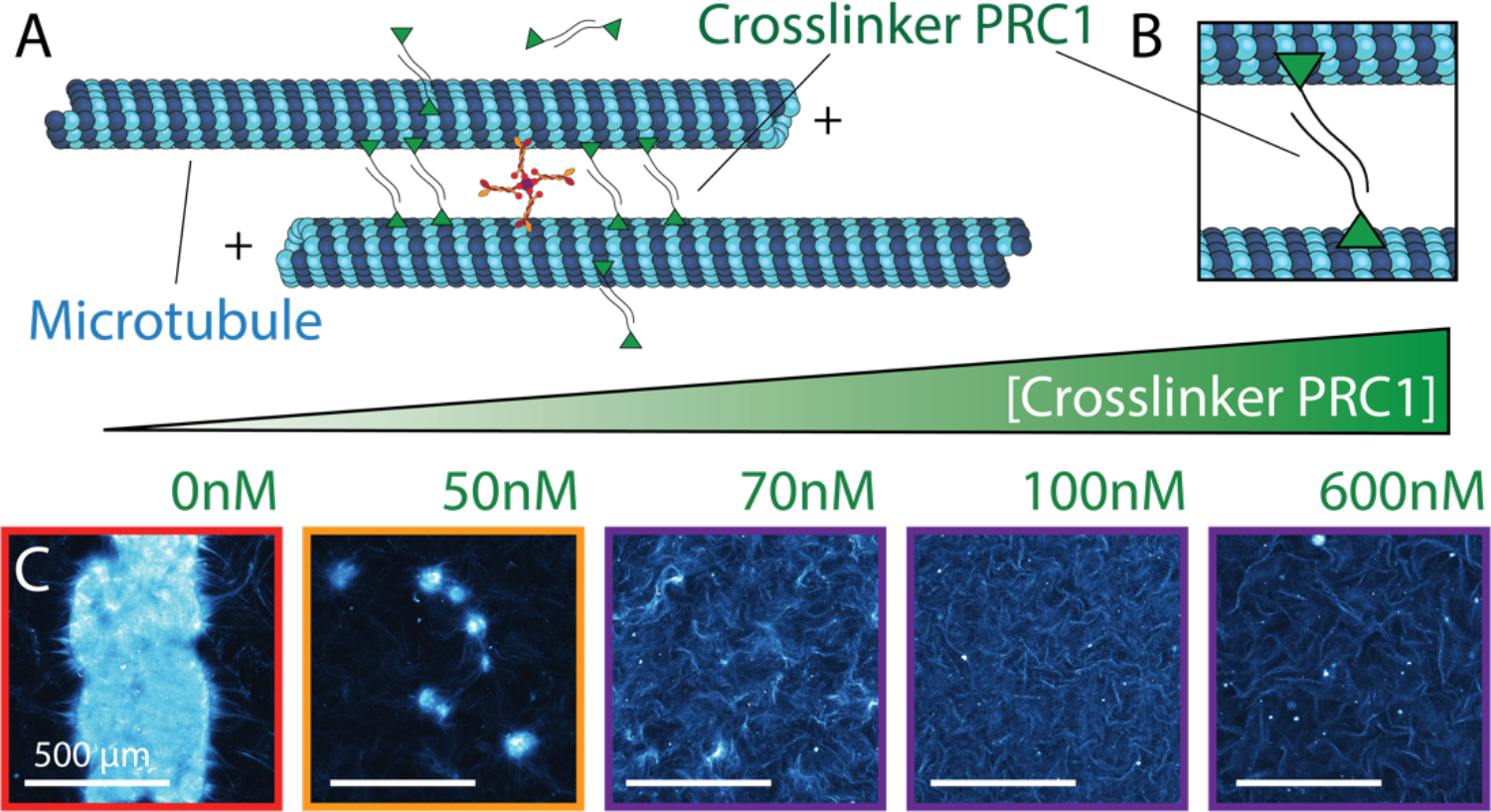
Adding PRC1 leads to a transition from contractile to extensile dynamics. A) Schematic of a microtubule bundle crosslinked by PRC1. B) Close up on a PRC1 dimer crosslinking two anti-parallel microtubules. C) Fluorescent pictures of a globally contractile phase with increasing concentrations of crosslinker (from 0 to 600nM). Adding an increasing amount of crosslinkers progressively transforms a globally contractile phase into an extensile phase. Images in C were taken with a confocal microscope. Scale bars are 500 µm. [ATP]=30 µM, [motor cluster]=480nM, [MT]=1.3mg/mL, [PEG]=0.6%

**Figure S8:**
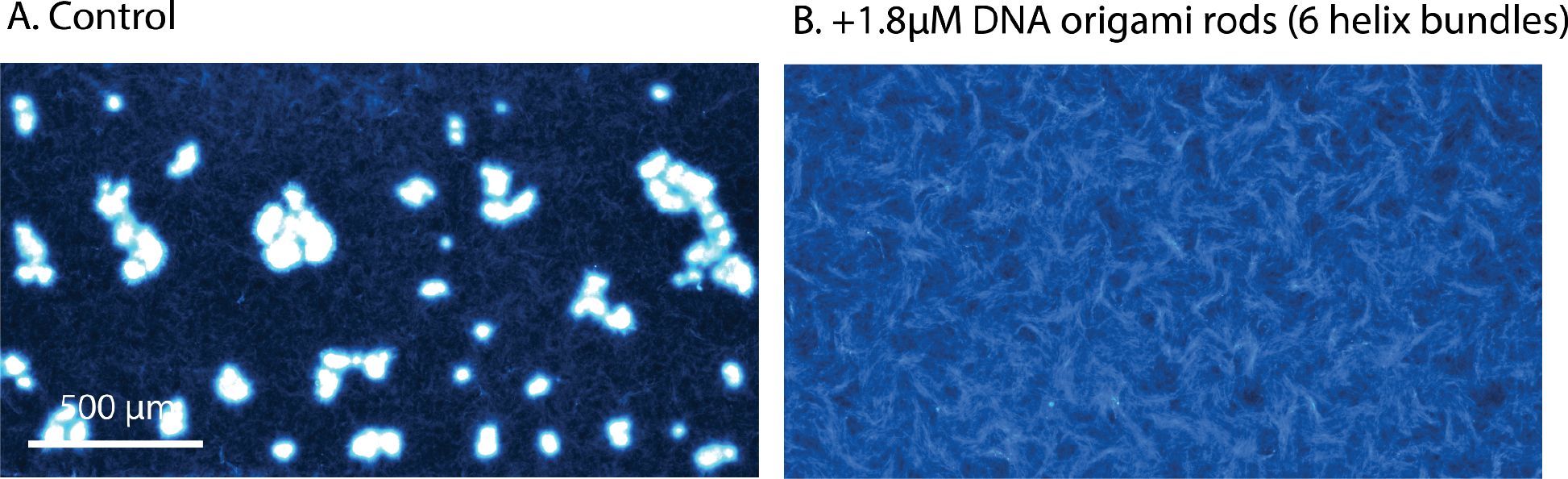
Adding 6-helix-bundles DNA origami rods turns a contractile phase extensile. A) Control is contractile, B) and turns extensile when 1.8uM of DNA rods are added. [ATP]=30µM, [Motor cluster]=400nM, [PRC1]=100nm, [MT]=1.3mg/mL

**Figure S9:**
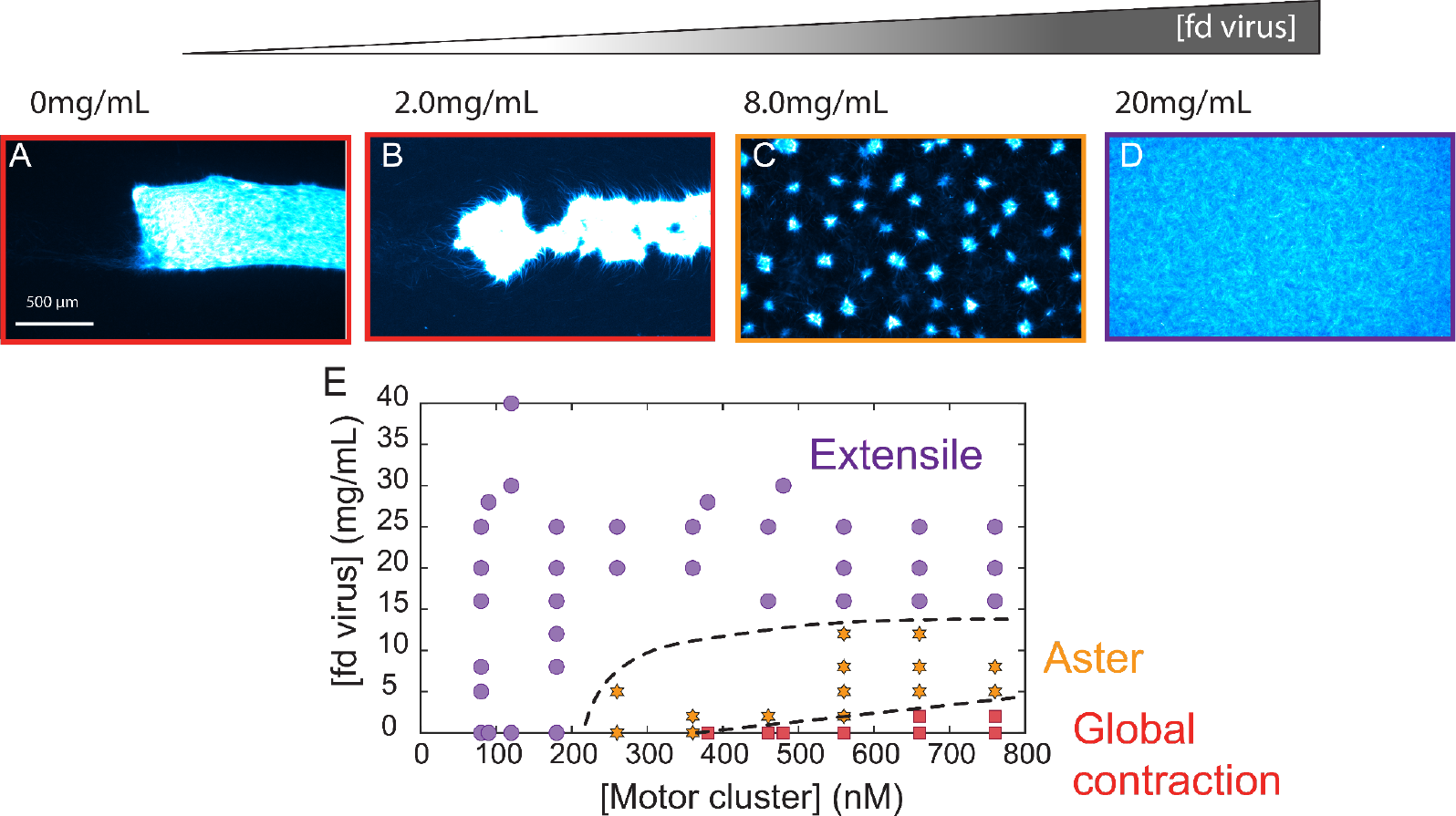
Effect of increasing nematic alignment by adding *fd* viruses to the active gel. A-D) fluorescent images when *fd* was added at 0, 2, 8, and 20 mg/mL. The globally contractile phase turns first into an aster phase, and then into the extensile phase when more *fd* viruses are added. E) *fd*-motor cluster phase diagram. Dash phase boundaries are a guide for the eye.

**Figure S10:**
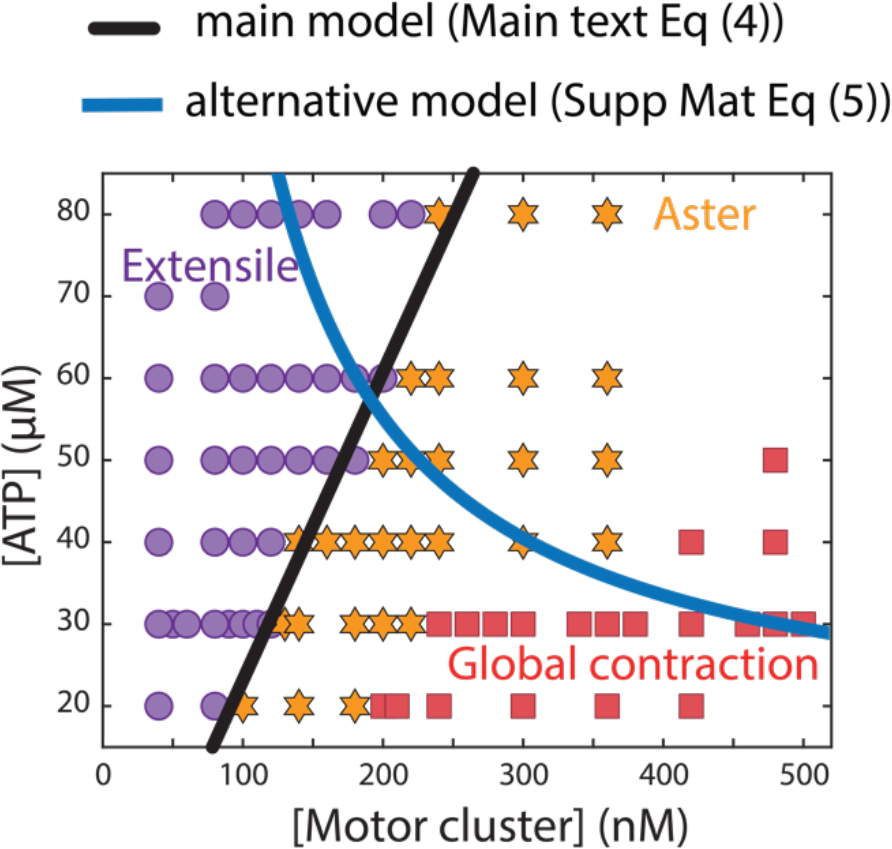
Alternative theoretical phase boundary overlayed on the experimental phase diagram shown in Fig. 1K. The blue continuous line is the alternative model where ATP-bound motor clusters contribute to end-adhesion (Supplementary Materials, section 5b, Supp Eq. 5). The black line is the model from the main text.

**Figure S11:**
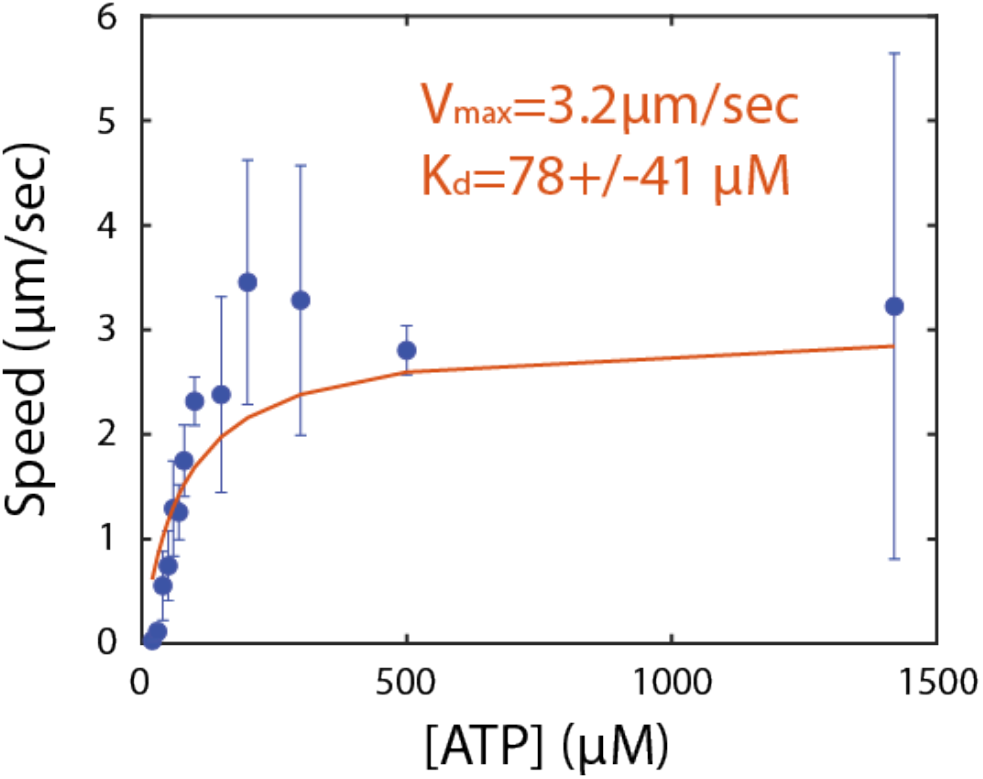
Estimation of K_d_. This plots shows how the average speed of the active extensile gels increases when ATP concentration increases. Error bars: standard deviation over 4 independent replicates. Blue points: experimental data points, red line: best fit of V=V_max_/(K_d_+[ATP]).

**Figure S12:**
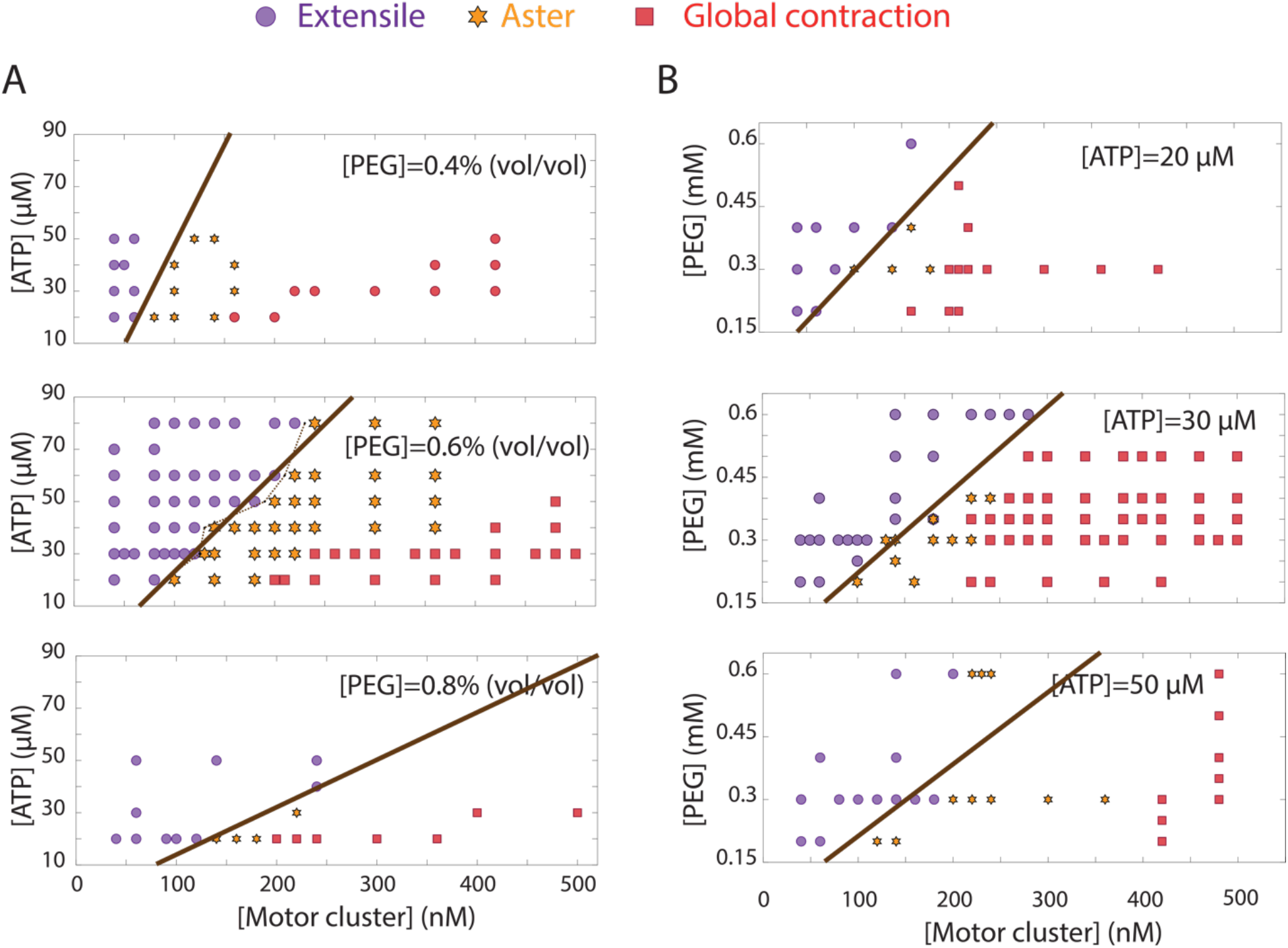
Summary of all the 2D phase diagrams collected for this study. A) ATP-motor cluster phase diagrams for various PEG concentrations. B) PEG-motor cluster phase diagrams for various ATP concentrations. [Tubulin]=1.33mg/mL for all experiments reported in these 6 phase diagrams. The dashed lines correspond to the experimental phase boundary to which the model (continuous line) is fitted.

**Figure S13:**
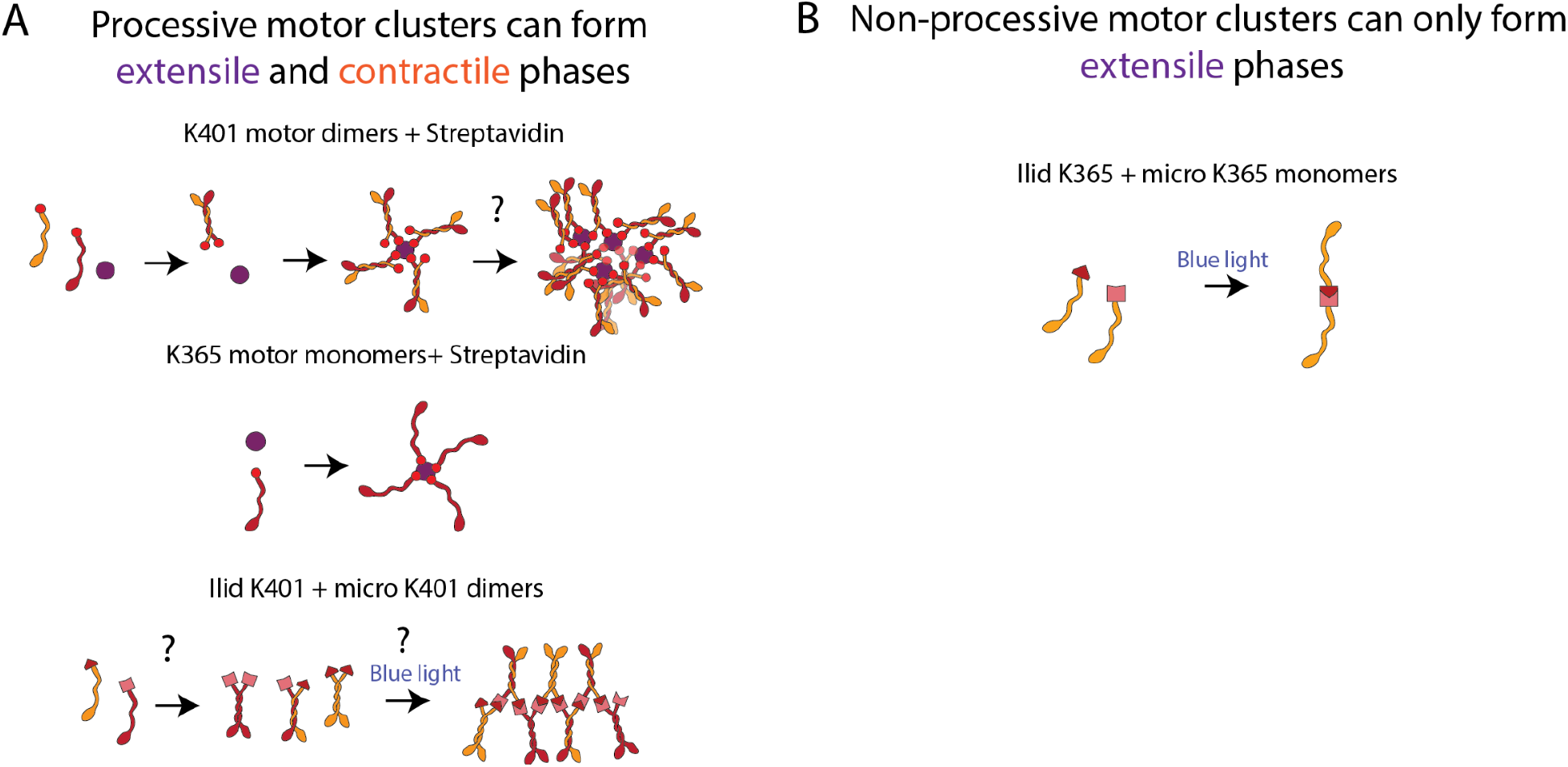
Valency of the motor clusters controls the self-organization of the kinesin-microtubule gels. A) multivalent clusters can drive extensile and contractile dynamics because the clusters are highly processive and have the ability to end-accumulate under the right experimental conditions. B) However, non-processive clusters cannot end-accumulate and therefore cannot form asters. These motors can only drive extensile dynamics.

### 7. Caption Supplementary videos

Video S1: extensile chaotic phase in an active network of microtubule and kinesin motor clusters. [ATP]=30µM, [motor cluster]=80nM, [PEG 20kDa]=0.6% vol/vol, microtubules labeled with Alexa 647.

Video S2: aster phase in an active network of microtubule and kinesin motor clusters. Asters locally contract while the background is still extensile. [ATP]=30µM, [motor cluster]=180nM, [PEG 20kDa]=0.6% vol/vol, microtubules labeled with Alexa 647.

Video S3: globally contractile phase in an active network of microtubule and kinesin motor clusters. [ATP]=30µM, [motor cluster]=380nM, [PEG 20kDa]=0.6% vol/vol, microtubules labeled with Alexa 647.

